# Direct ionic stress sensing and mitigation by the transcription factor NFAT5

**DOI:** 10.1101/2023.09.23.559074

**Authors:** Chandni B. Khandwala, Parijat Sarkar, H. Broder Schmidt, Mengxiao Ma, Maia Kinnebrew, Ganesh V. Pusapati, Bhaven B. Patel, Desiree Tillo, Andres M. Lebensohn, Rajat Rohatgi

**Affiliations:** Departments of Biochemistry and Medicine, Stanford University School of Medicine, Stanford, CA 94305, USA.; Laboratory of Cellular and Molecular Biology, Center for Cancer Research, National Cancer Institute, National Institutes of Health, NIH, Building 37, RM 2056B, Bethesda, MD, 20892, USA.; Centerfor Cancer Research Genomics Core, National Cancer Institute, National Institutes of Health, NIH, Building 37, RM 2056B, Bethesda, MD, 20892, USA.

**Keywords:** NFAT5, TonEBP, hypertonic stress, cell volume regulation, ion homeostasis, ionic strength, biomolecular condensates, phase separation, prions, ionic stress, transcription, osmotic stress, macromolecular crowding, osmolytes, intrinsically disordered protein, HOG pathway, tonicity

## Abstract

Homeostatic control of intracellular ionic strength is essential for protein, organelle and genome function, yet mechanisms that sense and enable adaptation to ionic stress remain poorly understood in animals. We find that the transcription factor NFAT5 directly senses solution ionic strength using a C-terminal intrinsically disordered region. Both in intact cells and in a purified system, NFAT5 forms dynamic, reversible biomolecular condensates in response to increasing ionic strength. This self-associative property, conserved from insects to mammals, allows NFAT5 to accumulate in the nucleus and activate genes that restore cellular ion content. Mutations that reduce condensation or those that promote aggregation both reduce NFAT5 activity, highlighting the importance of optimally tuned associative interactions. Remarkably, human NFAT5 alone is sufficient to reconstitute a mammalian transcriptional response to ionic or hypertonic stress in yeast. Thus NFAT5 is both the sensor and effector of a cell-autonomous ionic stress response pathway in animal cells.

## INTRODUCTION

The importance of water for the function of living organisms has been recognized since antiquity. Organisms across the tree of life have evolved molecular, cellular and organismal strategies to sense and adapt to water stress^1^. Water stress is a particular concern for terrestrial organisms that face fluctuating or unpredictable access to water. Understanding how our cells and tissues respond to water deprivation is imperative for human health in this era of accelerating climate change. The combination of rising temperatures (which increase evaporative water losses) and scarcity of potable water is contributing to the emergence and expansion of diseases (such as Chronic Kidney Disease of unknown origin) that may be caused by the effects of water stress on vulnerable tissues.^2^

At the level of individual cells, water stress manifests as an increase in the osmotic pressure gradient across the plasma membrane (hereafter referred to as “hypertonic stress”) (reviewed in^3–5^). Extracellular osmolarity rises above cytoplasmic osmolarity due to an increase in the concentration of charged ions and small, non-membrane permeable organic molecules. Thus, water stress can be caused by environmental water loss (e.g. desiccation, evaporation or freezing) or by an increase in the extracellular concentration of non-permeable solutes (e.g. rising salinity).

Cellular responses to hypertonic stress rely on the fundamental chemical principle that osmotic pressure is a colligative property, dependent on the collective number of solute particles rather than their specific identities. The acute consequence of hypertonic stress is cell shrinkage as water moves down its chemical potential gradient from inside to outside the cell. Both cell shrinkage and the adaptive mechanisms of cell volume recovery (called Regulatory Volume Increase or RVI) result in a rise in the cytoplasmic concentrations of charged ions.^5,6^ During the acute RVI response, the rise in extracellular osmolarity is balanced by a rise in the concentration of intracellular ions, allowing cytoplasmic rehydration and cell volume recovery on the timescale of minutes. However, this comes at the cost of an increase in the intracellular content of ions that is incompatible with long-term cell survival. An altered cytoplasmic ionic composition disrupts the structure and function of proteins, organelles and even the genome.^4,5,7,8^

Comparative physiologists have made the surprising observation that cytoplasmic ionic strength is conserved between organisms from all branches of life that live in markedly diverse habitats.^4,9^ A revealing exception is provided by certain halophilic bacteria that have high molar concentrations of cytoplasmic ions. The proteomes of these halobacteria have undergone extensive sequence changes during evolution to function in this extreme environment, but the consequence of this adaptive strategy is that they can only live in highly hyperosmotic habitats.^10^ For most other organisms, the preferred strategy has been the evolution of osmoregulatory systems that allow the maintenance of a stable intracellular ionic milieu despite variable extracellular osmolarity.^4^ This stable cytoplasmic ionic content allows protein function to be maintained in environments with fluctuating osmolarities without adaptive changes in protein sequence.

Osmoregulatory systems show a remarkable degree of convergent evolution across the tree of life. In organisms as diverse as plants, bacteria, fungi, fishes, invertebrates and mammals, the adaptation to hypertonic stress depends on the intracellular accumulation of small organic osmolytes.^4,9,11^ Osmolytes are non-membrane permeable molecules that are uncharged or zwitterionic and include polyhydric alcohols (e.g. glycerol, sorbitol, myo-inositol), free amino acids (e.g. taurine, proline, alanine) and methylamines (e.g. betaine). Most importantly, osmolytes have stabilizing or neutral effects on protein structure and activity across a wide range of concentrations. Indeed, osmolytes have structural similarities to protein stabilizing ions of the Hofmeister series and are often used to stabilize recombinant proteins in the laboratory.^4^ Thus, nearly all cell-autonomous strategies to adapt to hyperosmotic stress depend on varying the cytoplasmic concentration of organic osmolytes to balance extracellular osmotic pressure, preventing the depletion of cellular water. This strategy allows the intracellular ionic strength to be maintained within the narrow range required for protein function.

Osmoregulatory systems have been studied in diverse organisms, revealing both transcriptional and non-transcriptional strategies to control the synthesis and uptake of osmolytes. An exemplar eukaryotic system is the <u>h</u>igh-<u>o</u>smolarity <u>g</u>lycerol (HOG) pathway in the yeast *Saccharomyces cerevisiae*.^12,13^ In the HOG pathway, membrane proteins presumed to be sensors of hypertonic stress activate a cytoplasmic kinase cascade centered on the p38/Mitogen-activated kinase (MAPK) Hog1, ultimately leading to transcriptional activation of genes that promote the synthesis and accumulation of glycerol, the major osmolyte in *S.cerevisiae*. Multicellular organisms like humans have developed both behavioral responses (e.g. water seeking driven by thirst) and organ-level responses (e.g. water conservation in the kidney) to mitigate elevated plasma osmolarity. However, all our cells have also retained the universal cell-autonomous strategy of accumulating intracellular osmolytes. These responses are particularly important in tissues exposed to hypertonic stress, including the renal medulla, lymphoid tissues, and inflamed tissues.^14^ Interstitial fluid osmolarity in the renal medulla can rise to 1200 mOsm/L in humans and 4000 mOsm/L in mice, many times greater than plasma osmolarity (∼300 mOsm/L) and even greater than the osmolarity of seawater (1000 mOsm/L).^15^ The high interstitial osmolarity of the medulla allows the kidney to reabsorb free water from the renal filtrate and excrete urine that is more concentrated than plasma, a physiological function essential for the homeostatic regulation of plasma osmolarity. Consequently, cells in the renal medulla are exposed to extreme hypertonic stress, which is mitigated by the intracellular accumulation of osmolytes, as it is across all branches of life.^14,16^

Nearly three decades ago, the Rel-family transcription factor NFAT5 (also known as Tonicity Enhancer Binding Protein, or TonEBP) was identified as the master regulator of the response to hypertonic stress in mammals (**Figure 1A**).^17,18^ NFAT5 activates the transcription of many osmoadaptive genes, including those that promote the synthesis and accumulation of osmolytes and those that encode heat shock proteins to manage protein folding stress.^19–21^ Despite the central importance of NFAT5 in enabling tissue resilience to hypertonic stress, the mechanism (analogous to the iconic HOG pathway in *S.cerevisiae)* by which hypertonic stress signals are sensed and conveyed to NFAT5 remains to be identified in animals. Thus, we set out to answer two central mysteries in the mammalian osmoregulatory response: how is hypertonic stress sensed and how is this signal transmitted to NFAT5?

**Figure 1.**
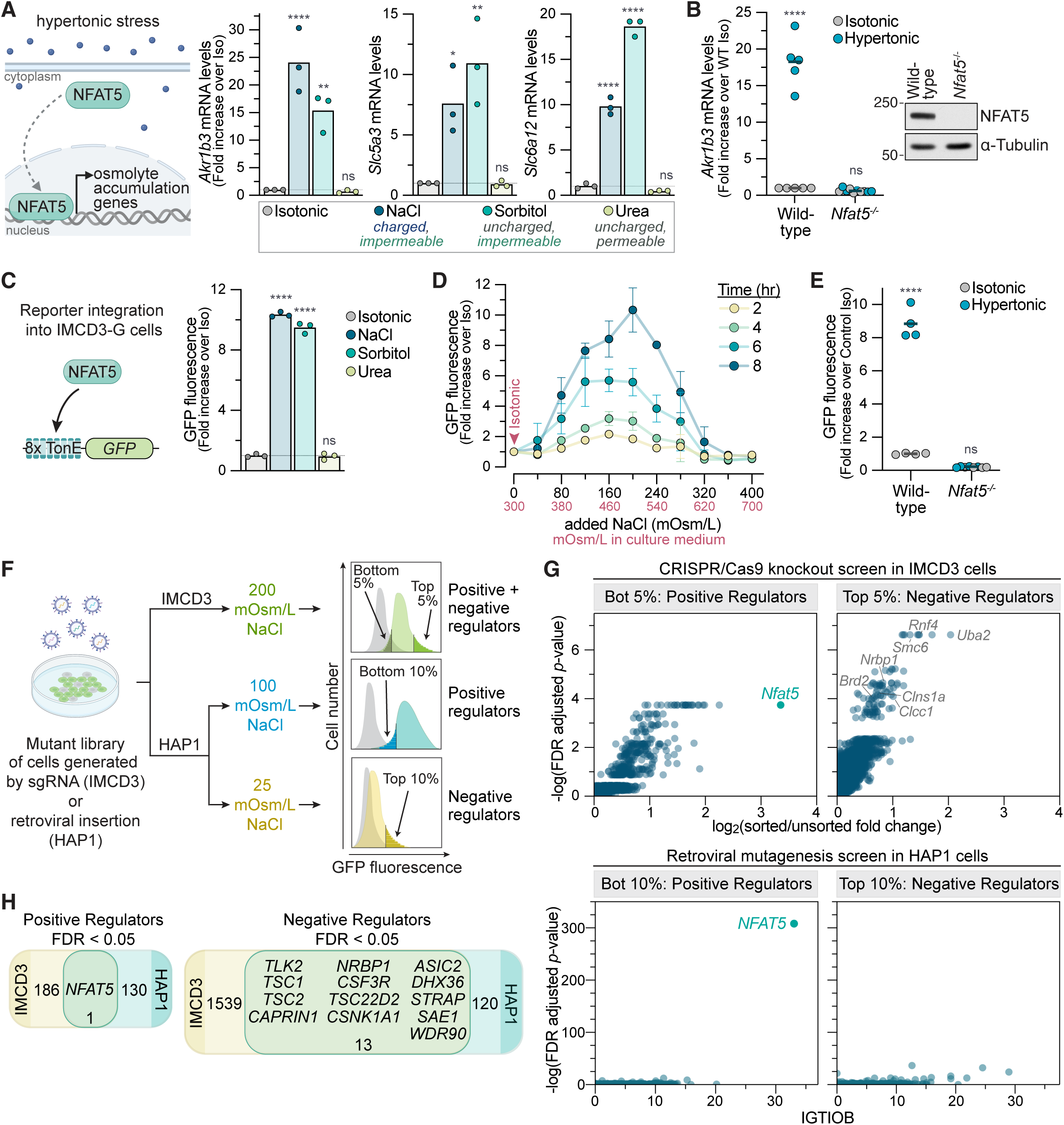
Genetic screens to identify positive and negative regulators of the transcriptional response to hypertonic stress. **(A)** The expression of three direct NFAT5 target genes was measured by quantitative reverse transcription PCR (RT-qPCR) after 8 hours (hrs) in isotonic media (300 mOsm/L) or after the addition of 200 mOsm/L of NaCl, sorbitol, or urea, raising the total media osmolarity in each case to ∼500 mOsm/L. **(B)** Expression of the NFAT5 target gene *Akr1b3* in wild-type (WT) IMCD3 cells or a clonal *Nfat5*^-/-^ cell line after 8 hrs in isotonic or hypertonic media (+200 mOsm/L NaCl). Elimination of NFAT5 in *Nfat5*^-/-^ cells was confirmed by immunoblotting (right). Data from four additional independent *Nfat5*^-/-^ clonal cell lines is shown in **Figure S1A**. **(C)** GFP fluorescence was measured by flow cytometry in IMCD3-G cells, which stably carry the 8TonE-GFP reporter (left), after 8 hrs in isotonic or hypertonic media (+200 mOsm/L NaCl, Sorbitol or Urea). Each point depicts the median GFP fluorescence from >2000 cells. **(D)** 8TonE-GFP reporter activity in IMCD3-G cells was measured by flow cytometry at the indicated time points in response to increasing amounts of NaCl added to isotonic media. Total media osmolarity at each concentration of NaCl is shown in red on the secondary x-axis. Each point shows the mean ± Standard Deviation (SD) of three independent median measurements, each from a population of >2000 cells. **(E)** 8TonE-GFP reporter activity after exposure to hypertonic media (+200 mOsm/L NaCl, 8 hrs) in WT or *Nfat5*^-/-^ IMCD3 cells. **(F)** The strategy used for genome-wide loss-of-function screens in mouse IMCD3 and human haploid (HAP1) cells using a stably integrated 8TonE-GFP reporter. Fluorescence activated cell sorting (FACS) was used to collect cells carrying mutations in genes encoding positive or negative regulators of the NFAT5 transcriptional response following exposure to hypertonic stress. The retroviral strategy used for insertional mutagenesis in HAP1 cells is shown in **Figure S1H**. **(G)** Volcano plots showing results from the genetic screens outlined in **F**. For the CRISPR screen (top) in IMCD3 cells, the x-axis shows the enrichment of each gene (calculated as the mean of all four sgRNAs targeting the gene) in the sorted population relative to the unsorted population and the y-axis shows statistical significance as measured by the false discovery rate (FDR)-corrected *p*-value. For the haploid screen (bottom) in HAP1 cells, the x-axis shows the Intronic Gene-trap Insertion Orientation Bias (IGTIOB) score, which scores the bias towards inactivating insertions in each gene from sorted cells only, and the y-axis shows the FDR-corrected *p*-value, reflecting the enrichment of gene-trap insertions in sorted over unsorted cells. **(H)** Venn diagrams representing overlapping positive and negative regulators with FDR-corrected *p*-values < 0.05 from the independent screens conducted in IMCD3 and HAP1 cells. **Statistics**: Bars (**A**,**C**) or black horizontal lines (**B**,**E**) denote mean values calculated from independent measurements shown as points. Statistical significance (**A**,**B**,**C**,**E**) was determined by a two-way ANOVA test with Sidak’s multiple comparison post-test (*n*>3 independent experiments). *p*-values symbol are: **** <0.0001, *** <0.001, **<0.01, and *<0.05. **See also Figure S1**.

**Supplementary Figure 1.**
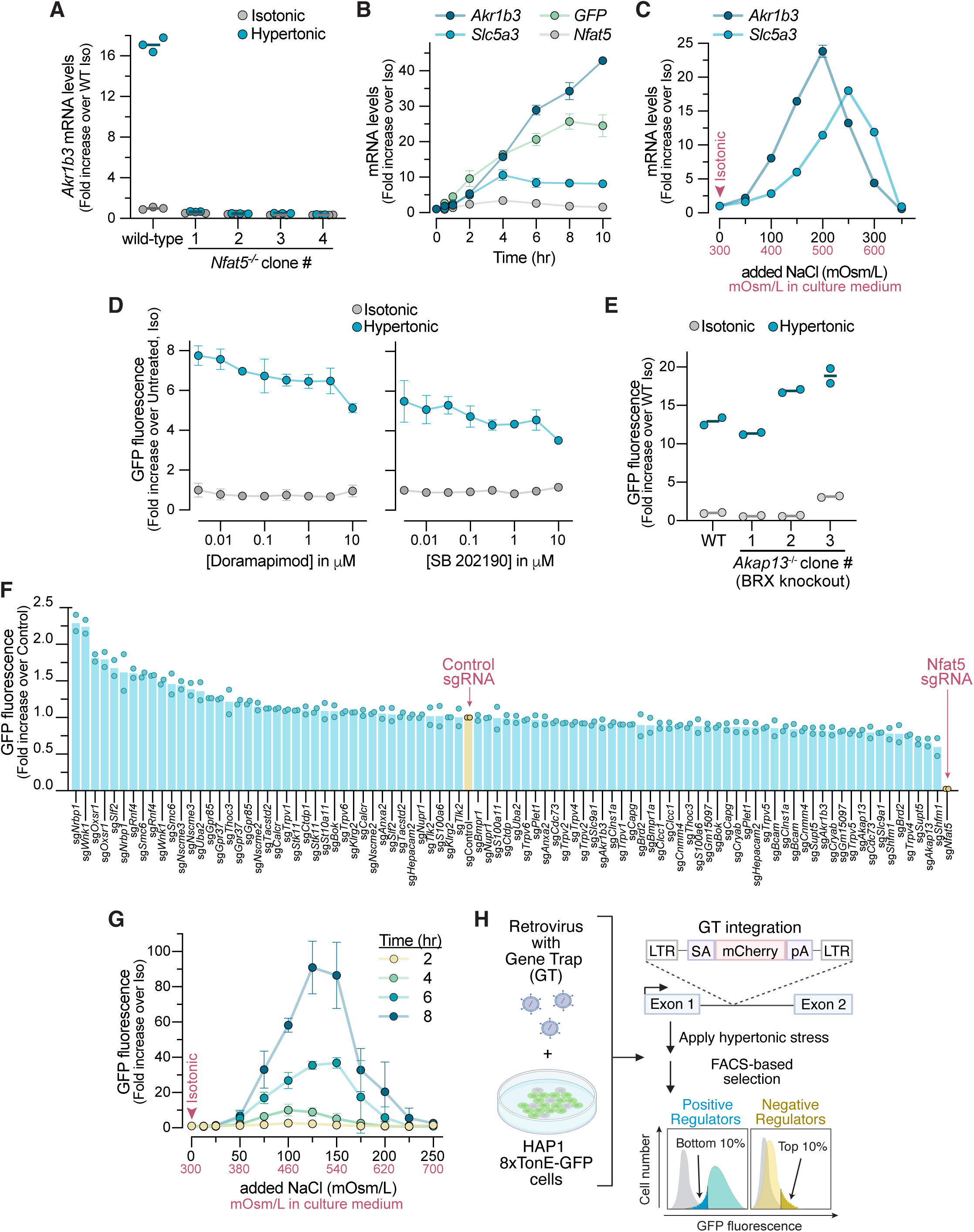
Optimization and validation of genetic screens for NFAT5 regulators, Related to Figure 1. **(A)** NFAT5 target gene expression measured by RT-qPCR in wild-type (WT) IMCD3 cells and four independent *Nfat5*^-/-^ clonal cell lines after 8 hrs in isotonic media (300 mOsm/L) or in hypertonic media (+200 mOsm/L NaCl, raising total media osmolarity to 500 mOsm/L). Black horizontal lines denote mean values calculated from 3 independent measurements shown as points. **(B)** Abundance of *Akr1b3*, *Slc5a3*, *Nfat5,* and the *GFP* reporter mRNA was measured using RT-qPCR in IMCD3-G cells after exposure to hypertonic stress (+200 mOsm/L NaCl) for the indicated time periods. Each point shows the mean ± Standard Deviation (SD) of three independent measurements. **(C)** Expression of *Akr1b3* and *Slc5a3* in WT IMCD3 cells treated with increasing amounts of NaCl added to isotonic media for 8 hrs. Total media osmolarity at each concentration of NaCl is shown in red on the secondary x-axis. Each point shows the mean ± SD of three independent measurements. **(D)** 8TonE-GFP reporter activity in IMCD3-G cells measured by flow cytometry after 8 hrs in isotonic or hypertonic (+200 mOsm/L NaCl) media in the presence of increasing concentrations of the p38 inhibitors doramapimod or SB 202190. Each point shows the mean ± SD of three independent median measurements, each from a population of >2000 cells. **(E)** 8TonE-GFP reporter activity measured by flow cytometry in WT IMCD3-G cells or three independent *Akap13*^-/-^ IMCD3-G clonal cell lines measured after 8 hrs in isotonic or hypertonic (+200 mOsm/L NaCl) media. Each point shows a median measurement from a population of >2000 cells; black horizontal lines show the mean of these independent median values. **(F)** 8TonE-GFP reporter activity after exposure to hypertonic media (+200 mOsm/L NaCl, 8 hrs) in IMCD3-G cells expressing Cas9 and a non-targeting control sgRNA or sgRNAs targeting the indicated genes on the x-axis. Each point shows an independent median measurement from a population of >2000 cells. **(G)** 8TonE-GFP reporter activity in human haploid (HAP1) cells at the indicated time points in response to increasing amounts of NaCl added to isotonic media. Each point shows the mean ± SD of three independent median measurements, each from a population of >2000 cells. Total media osmolarity at each concentration of NaCl is shown in red on the secondary x-axis. **(H)** Strategy used for the retroviral insertional mutagenesis screen in human HAP1 cells using the stably integrated 8TonE-GFP reporter. Insertion of the gene-trap (GT) cassette, positioned between the retroviral long terminal repeats (LTRs), in the sense orientation into an intron will trigger transcriptional termination: the splice acceptor (SA) will cause this cassette to be spliced to the preceding exon, leading to transcriptional termination due to a strong polyadenylation (pA) signal. Fluorescence activated cell sorting (FACS) was used to collect cells carrying mutations in genes encoding positive or negative regulators of the NFAT5 transcriptional response following exposure to hypertonic stress (see **Figure 1F**).

## RESULTS

### Genetic screens to identify regulators of NFAT5

Genome-wide screens in cultured cells are powerful and unbiased discovery tools to identify genes that act cell-autonomously to regulate a signaling pathway. We have previously relied on both CRISPR/Cas9-based methods and insertional mutagenesis-based screens to identify new components in the Hedgehog and WNT signaling pathways using fluorescence-based transcriptional reporters for phenotypic enrichment.^22,23^ To apply our CRISPR/Cas9-based screening pipeline to investigate the NFAT5 pathway, we chose mouse Inner Medullary Collecting Duct (IMCD3) cells, a cell line derived from the renal medulla that has been extensively used to study cellular responses to hypertonic stress.^24^ Increasing the osmolarity of IMCD3 culture media by adding charged (NaCl) or uncharged (sorbitol) molecules that are membrane impermeable increased the expression of three established NFAT5 target genes that promote intracellular osmolyte accumulation: *Akr1b3* (aldol reductase, required for sorbitol synthesis), *Slc5a3* (a myo-inositol importer) and *Slc6a12* (a betaine importer) (**Figure 1A**).^20^ As expected, the disruption of *Nfat5* abrogated these transcriptional responses (**Figure 1B** and **S1A**). As a control, we added urea to IMCD3 culture media. Urea, which is also enriched in the renal medulla, is a membrane permeable small molecule that raises the intracellular and extracellular osmolarity equally.^19^ While urea increases overall osmolarity, it does not impose an osmotic pressure gradient or drive water movement across the plasma membrane and, consequently, failed to increase *Akr1b3*, *Slc5a3* or *Slc6a12* mRNA abundance (**Figure 1A**).

We engineered a clonal IMCD3 reporter cell line (IMCD3-G) stably expressing a synthetic reporter for NFAT5 activity, which we call 8TonE-GFP (**Figure 1C**). In this reporter, the expression of Green Fluorescent Protein (GFP) is driven by a tandem array of eight Tonicity Enhancer (TonE) DNA binding sites for NFAT5.^25^ GFP fluorescence in IMCD3-G cells increased specifically in response to hypertonic stress in a dose-, time-and NFAT5-dependent manner (**Figure 1D** and **1E**). The kinetics of GFP mRNA accumulation resembled those of endogenous NFAT5 target genes (**Figure S1B**). As noted previously^26^ but not further investigated in our current study, increasing hypertonic stress resulted in a bell-shaped change in NFAT5 activity, measured either using 8TonE-GFP fluorescence (**Figure 1D**) or endogenous target gene mRNA abundances (**Figure S1C**). Most of our experiments were conducted using conditions in the ascending phase of this dose-response curve.

We conducted loss-of-function CRISPR screens designed to find positive and negative regulators of NFAT5 activity in IMCD3-G cells using the genome-wide Brie library (**Figure 1F**).^27^ *Nfat5* itself was the top hit from the screen for positive regulators (**Figure 1G**). Many of the other components previously implicated in NFAT5 activation (e.g. BRX, p38/MAPK) failed to score as statistically significant hits (**Figures S1D** and **S1E**).^28^ We validated 50 of the top screen hits using independent single guide RNAs (sgRNAs) in IMCD-G cells, comparing the effects of each candidate gene’s disruption to that of NFAT5 itself (**Figure S1F**). Based on prior experience with Hh and WNT pathway screens, we expected the disruption of any gene encoding a core component of the NFAT5 activation pathway to have an effect size commensurate with the disruption of *Nfat5* itself. However, targeted disruption of most of the genes identified in the screen had a much smaller effect on 8TonE-GFP activation, suggesting that our CRISPR screen had identified modulators, but had failed to identify any obligate components of the NFAT5 pathway (**Figure S1F**).

Given the unexpected results of our CRISPR screen in mouse IMCD3 cells, we performed a second set of genetic screens in human haploid cells (HAP1 cells) using insertional mutagenesis with a retrovirus to generate the genome-wide mutant library (**Figures S1G** and **S1H**).^29^ We used the same hypertonicity-responsive 8TonE-GFP reporter to select for positive and negative regulators of NFAT5 activity in two separate screens (**Figure 1F**). Remarkably, *NFAT5* was the most statistically significant hit in this haploid screen (**Figure 1G**). It was also the only common positive regulator identified in both the CRISPR and haploid screens (**Figure 1H**). Much to our surprise, genetic screens in cell lines from two different species using two different mutagenesis methods identified NFAT5 as the only component required for the activation of a transcriptional reporter of hypertonic stress. We note that for both our screens, as for all forward genetics screening approaches, redundant and essential genes would not be expected to score as hits.

### Reconstitution of NFAT5 activity in *S. cerevisiae*

Our genetic screens led us to consider the hypothesis that NFAT5 alone may be sufficient to mediate and activate the transcriptional response to hypertonic stress in mammals and that no elaborate upstream signaling pathway analogous to the yeast HOG pathway is involved. A conclusive demonstration of this hypothesis would be that NFAT5 should be the only component required to endow a unicellular organism lacking *Nfat* genes with a mammalian-like hypertonic stress response. We chose *S. cerevisiae* for several reasons. First, fungi and animals diverged nearly 1.5 billion years ago, and their osmoregulatory pathways seem to be quite different.^30^ *S. cerevisiae* lacks an *Nfat5*-related gene. In addition, *S. cerevisiae* has a cell wall (which is not present in mammalian cells) and is thought to sense hypertonic pressure through changes in turgor pressure, using membrane protein sensors that are not conserved in mammals.^31^ Finally, the human *NFAT5* cDNA originally cloned using a one-hybrid screen, showing that NFAT5 can activate the yeast transcriptional machinery.^17^

The expression of full-length NFAT5, which is ∼1500 amino acids (a.a.) in length, was challenging in *S. cerevisiae.* Instead, we engineered a minimal NFAT5 (hereafter “mini-NFAT5”) that could support hypertonicity-regulated gene expression in IMCD3 cells but was compact enough to express in yeast (see **Figures 2A** and **S2A** for the domain structure of NFAT5 and variants). NFAT5 variants were tested for their ability to restore hypertonicity-induced target gene expression in *Nfat5*^-/-^ IMCD3 cells when stably expressed at moderate levels from a single genomic locus using Flp-mediated recombination (**Figure S2B**). While NFAT5 accumulates in the nucleus in response to hypertonic stress (**Figure 2B**), nuclear localization alone did not drive activation of target genes: an NFAT5 variant constitutively localized in the nucleus by fusion to a strong nuclear localization sequence (NLS) was properly regulated by hypertonic stress (**Figure S2C** and **S2D**). Thus, the cellular mechanism that detects and transduces the hypertonic stress signal can activate NFAT5 even when it is confined to the nucleus. We also took advantage of a previous truncation analysis of NFAT5 that identified two segments within its C-terminal domain (CTD) that were sufficient to confer hypertonicity-activated gene transcription when fused to a heterologous DNA binding domain.^32^ Combining these two insights, we engineered a mini-NFAT5 that contained a fluorescent protein for easy detection, an NLS for constitutive nuclear localization, the native Rel-homology DNA binding domain (DBD), and a portion of the native CTD (**Figure 2A**). Mini-NFAT5 was sufficient to drive activation of endogenous NFAT5 target genes in response to hypertonic stress, albeit to a lesser extent than the full-length protein (**Figure 2C**).

To test the activity of mini-NFAT5 in *S.cerevisiae*, we stably expressed (1) a variant of the 8TonE-GFP reporter constructed from a minimal yeast promoter (8TonE-*pCYC1*-GFP) and (2) mRuby3 tagged mini-NFAT5, codon optimized for yeast expression, driven by a galactose-inducible promoter (**Figure 2D**). *S.cerevisiae* strains expressing both transgenes were treated with galactose to induce expression of mini-NFAT5 (**Figure S2E**) and then subjected to increasing levels of hypertonic stress by the addition of NaCl to culture media. In multiple independent clonal strains, NaCl induced the dose-dependent activation of the 8TonE-*pCYC1-* GFP reporter as measured by GFP fluorescence (**Figure 2E**), GFP mRNA abundance (**Figure S2F**), or GFP protein abundance (**Figure S2G**). Consistent with the behavior of NFAT5 target genes in mammalian cells, reporter activation was induced only by non-membrane permeable solutes (NaCl, sorbitol) that exert an osmotic pressure gradient across the plasma membrane, but not by membrane permeable molecules like urea (**Figure 2F**, compare to **Figure 1A**).

**Figure 2.**
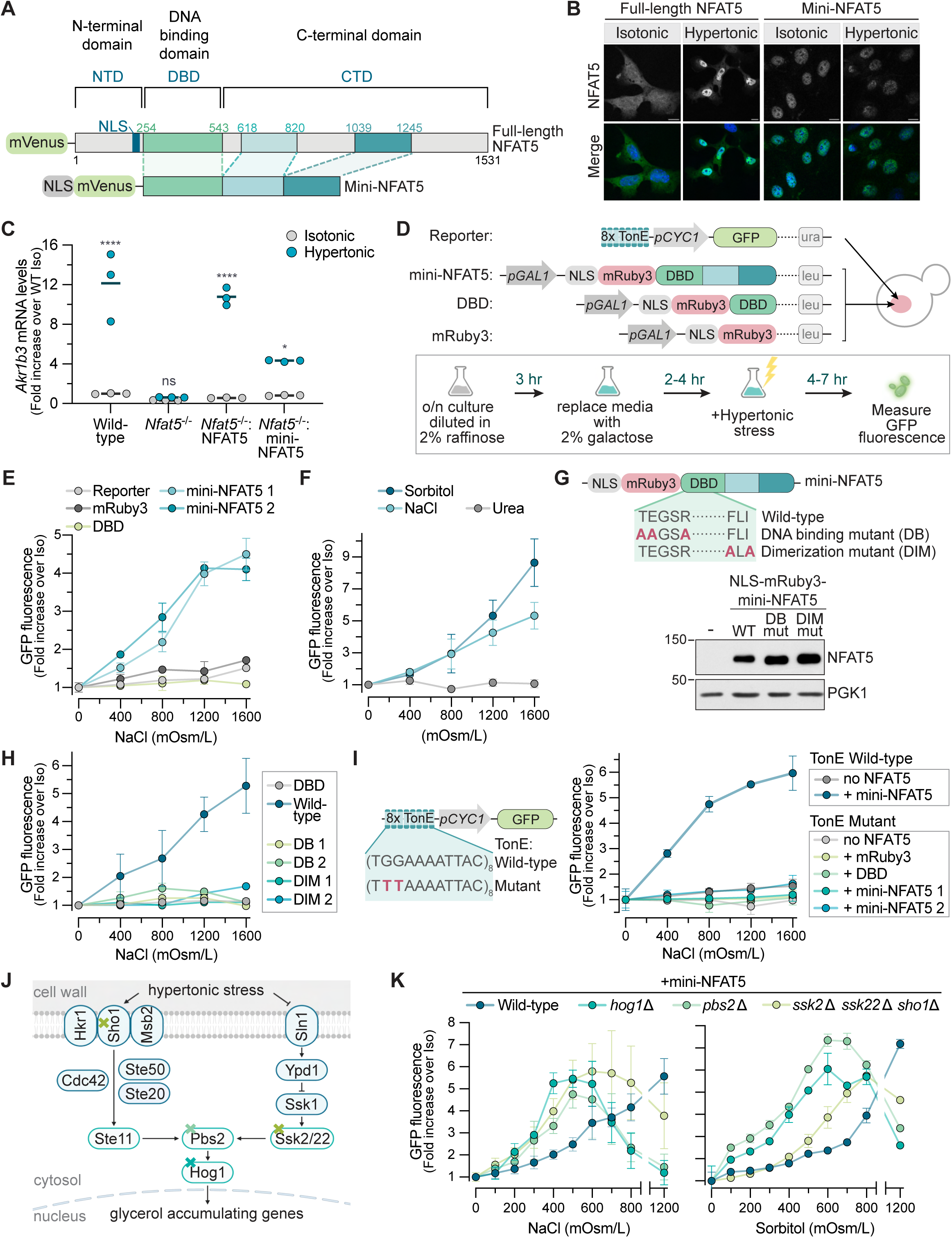
NFAT5 can be activated by hypertonic stress in *Saccharomyces cerevisiae.* **(A)** Domain structures of mVenus-tagged human full-length and mini-NFAT5. Mini-NFAT5 was engineered by joining two regions within the C-terminal domain (CTD) of NFAT5 to its DNA Binding Domain (DBD) and a strong heterologous nuclear localization sequence (NLS). **(B)** The subcellular localization of mVenus-tagged full-length or mini-NFAT5 stably expressed in IMCD3 cells exposed to isotonic or hypertonic (+200 mOsm/L NaCl) media for 30 minutes (min). Fluorescence signal from the mVenus tag fused to NFAT5 is shown alone (top) or merged with a DNA stain for nuclei (DAPI, bottom). Scale bars: 10 μm. **(C)** Expression of an NFAT5 target gene in WT cells, *Nfat5^-/-^* cells, or *Nfat5^-/-^* IMCD3 cells stably expressing either full-length or mini-NFAT5 (see **A**) after 8 hrs in isotonic or hypertonic (+200 mOsm/L NaCl) media. **(D)** Structure of the 8TonE-*pCYC1*-GFP reporter and galactose-inducible mini-NFAT5 variant genes integrated into wild-type W303 yeast cells (top). Schematic of the experimental workflow used to test reporter activity by flow cytometry in yeast cells following galactose (Gal) induction of the NFAT5 variants and hypertonic stress (bottom). **(E)** 8TonE-*pCYC1*-GFP reporter activity in yeast cells expressing the reporter only or the reporter in addition to mRuby3, mRuby3-DBD, or mRuby3-mini-NFAT5 (2 independent clones) in response to increasing amounts of NaCl added to complete synthetic media (CSM). **(F)** 8TonE-*pCYC1*-GFP reporter activity in yeast cells expressing mini-NFAT5 in response to increasing concentrations of NaCl, sorbitol, or urea. **(G)** Two separate sets of mutations introduced in the DBD in NLS-mRuby3-mini-NFAT5 to abrogate DNA-binding (DB) or dimerization (DIM). Abundances of the proteins encoded by these mini-NFAT5 variant constructs were compared by immunoblotting (bottom). **(H)** 8TonE-*pCYC1*-GFP reporter activity in yeast cells expressing the indicated variants of mini-NFAT5 exposed to increasing amounts of NaCl for 4 hrs. Two independent clones of the DNA-binding (DB) and dimerization (DIM) mutants (see **G**) were tested. **(I)** Sequence of the WT tonicity enhancer (TonE) binding site, compared to a mutant version known to abrogate its interaction with NFAT5. Graph on the right shows activity of the WT or mutant 8TonE-*pCYC1-* GFP reporter in yeast cells expressing the indicated variants of mini-NFAT5 in response to increasing concentrations of NaCl. **(J)** The HOG (high-osmolarity glycerol) pathway in *S. cerevisiae*. Coloured X’s denote three different genes or gene sets that were deleted to disrupt the pathway at various levels: *HOG1*, *PBS2*, or the combined triple deletion of *SSK2*, *SSK22* and *SHO1*. **(K)** 8TonE-*pCYCI*-GFP reporter activity in WT, *hogl*△, *pbs2*△, or *ssk2& ssk22*△ *shol*△ cells (also expressing mini-NFAT5) in response to increasing amounts of NaCl (left) or sorbitol (right). **Statistics**: Each point (**E**,**F**,**H**,**I**,**K**) shows the mean ± SD of >3 independent median measurements, each from a population of >5000 cells. Solid horizontal lines (**C**) denote mean values calculated from three independent measurements shown as points. Statistical significance (**C**) was determined by a two-way ANOVA test with Sidak’s multiple comparison post-test (*n*>3 independent experiments). P-value symbols are: **** *p*- value<0.0001 and * *p*-value<0.05. **See also Figure S2**.

**Supplementary Figure 2.**
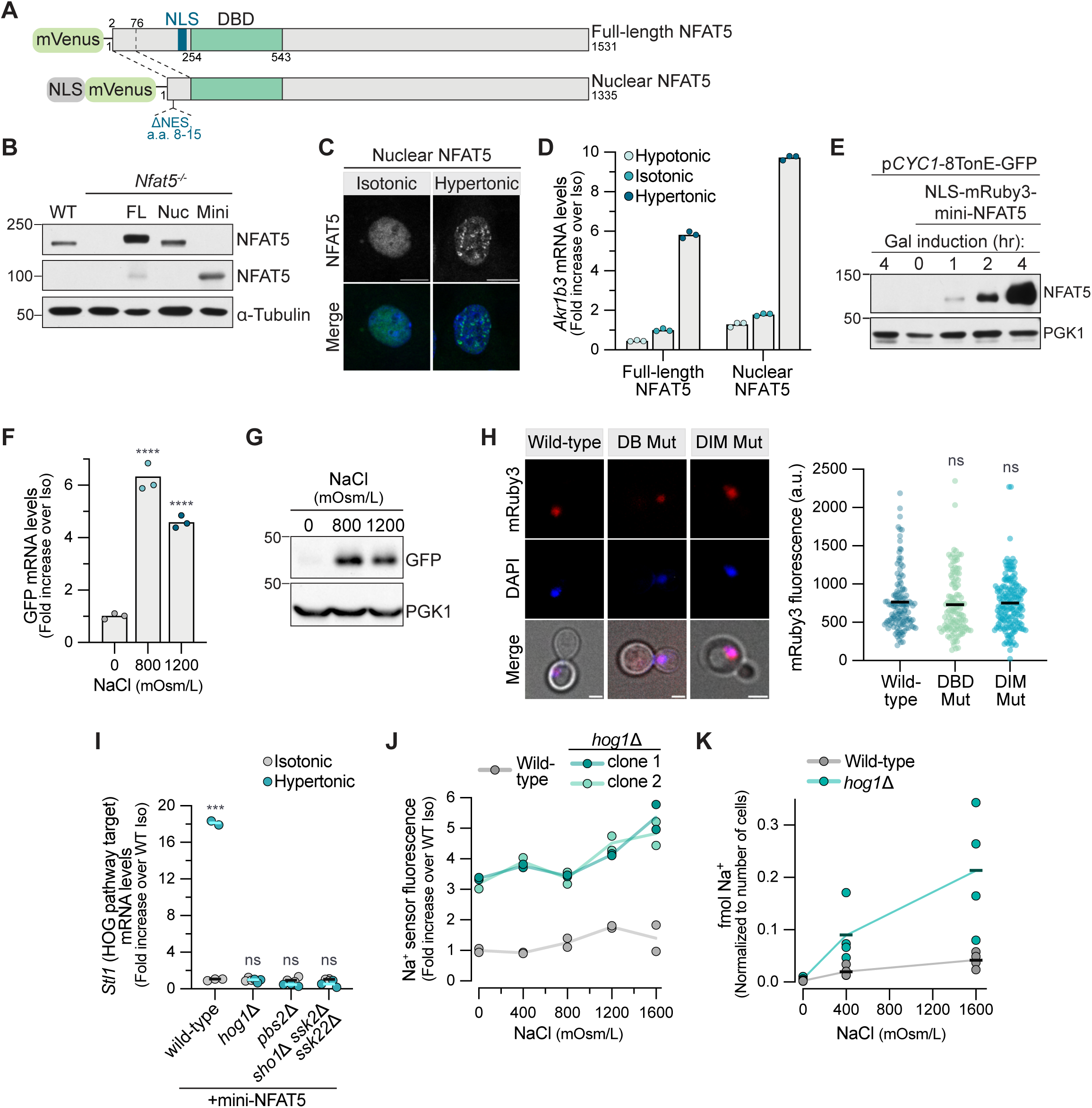
Characterization of mini-NFAT5 in IMCD3 and yeast cells, Related to Figure 2. **(A)** Diagram of full-length, mVenus-tagged NFAT5 (top) compared to a variant (Nuclear NFAT5) constitutively targeted to the nucleus by removal of its endogenous nuclear import and export sequences (a.a. 77-253) and addition of a heterologous strong nuclear localization sequence (NLS). **(B)** Immunoblot comparing NFAT5 abundance in WT, *Nfat5*^-/-^, or *Nfat5*^-/-^ IMCD3 cells stably expressing full­length (FL), nuclear (nuc), or mini-NFAT5. **(C)** Subcellular localization of nuclear NFAT5 after 30 min in isotonic or hypertonic (+200 mOsm/L NaCl) media. Fluorescence signal from the mVenus tag fused to NFAT5 is shown alone (top) and merged with a DNA stain (DAPI) to mark nuclei (bottom). Scale bar: 10 μm. **(D)** Expression of an NFAT5 target gene in *Nfat5*^-/-^ cells stably expressing full-length or nuclear NFAT5 after 8 hrs in hypotonic, isotonic, or hypertonic media (+200 mOsm/L NaCl). **(E)** Abundance of NLS-mRuby3-mini-NFAT5 at various times after galactose addition in a yeast strain used to test activation of NFAT5 by hypertonic stress. This strain contains two stably integrated transgenes, one encoding NLS-mRuby3-mini-NFAT5 driven by a galactose-inducible promoter and a second encoding the 8TonE-*pCYC1*-GFP NFAT5 reporter (see **Figure 2D**). The leftmost lane shows a control strain lacking the mini-NFAT5 transgene but containing the reporter. **(F)** Abundance of GFP mRNA in yeast cells expressing the 8TonE-*pCYC1*-GFP reporter and NLS-mRuby3-mini-NFAT5 was measured by RT-qPCR after 2 hrs in CSM containing the indicated concentrations of NaCl. **(G)** Abundance of GFP protein in yeast cells expressing the 8TonE-*pCYC1*-GFP reporter and NLS-mRuby3-mini-NFAT5 was measured by immunoblotting after 4 hrs in CSM containing the indicated concentrations of NaCl. **(H)** Representative images (left) showing subcellular localization of wild-type NLS-mRuby3-mini-NFAT5 and variants carrying mutations that abrogate DNA binding (DB Mut) or dimerization (DIM Mut) (see **Figure 2G**) in yeast cells used in the experiment depicted in **Figure 2H**. Graph on right shows quantitation of nuclear mRuby3 fluorescence in single yeast cells (measured from images of the type shown on the left) expressing the indicated NFAT5 variant (*n*>118 cells per strain). **(I)** Expression of the HOG pathway target gene *Stl1* in wild-type, *hogl*△, *pbs2*△, or *ssk2A ssk22*△ *shol*△ cells after 2 hrs in CSM supplemented with 1200 mOsm/L NaCl. **(J)** Fluorescence of a sodium sensor (see methods) in wild-type and *hogl*△ cells (2 independent clones) in response to increasing amounts of NaCl added to CSM. Two technical replicates were measured for each clone and are shown separately. **(K)** Abundance of sodium ions was measured by inductively coupled plasma optical emission spectroscopy (ICP-OES) in wild-type and *hogl*△ cells. **Statistics**: Bars (**D**,**F**) or horizontal lines (**H**,**I,K**) denote mean values calculated from independent measurements shown as points. Statistical significance was determined by a one-way (**F**) or two-way (**I**) ANOVA test with Sidak’s multiple comparison post-test (*n*>3 independent experiments) or (**H**) by the Kruskal Wallis test. *p*-values symbols are: **** <0.0001 and *** <0.001.

We confirmed the specificity of the *S.cerevisiae* mini-NFAT5 response in several ways. First, the transactivation domain from the NFAT5 CTD was required for activity: mRuby3 alone or mRuby3 fused to the isolated DBD of NFAT5 were both inactive (**Figure 2E**). Previous structural and biochemical studies have identified point mutations in the NFAT5 DBD that abolish activity in mouse or human cells by either preventing DNA binding or dimerization (**Figure 2G**).^33^ Both classes of mutations abolished the ability of mini-NFAT5 to activate the 8TonE-*pCYC1*-GFP reporter in *S.cerevisiae* (**Figure 2H**) without changing protein abundance or nuclear localization (Figures 2G and **S2H**). Finally, mutations in the TonE binding sites of the 8TonE-*pCYC1*- GFP reporter known to abolish NFAT5 binding in mammalian systems also abrogated the mini-NFAT5 transcriptional response to hypertonic stress (**Figure 2I**).^34^ Taken together, these results show that the sequence requirements for NFAT5 activity are similar in both *S.cerevisiae* and mammalian systems, supporting a shared mechanism of transcriptional activation in response to hypertonic stress.

In our studies, hypertonic stress applied to *S.cerevisiae* would also be predicted to activate the HOG pathway. Thus, it was important to test whether HOG signaling mediates the activation of mammalian mini-NFAT5 expressed in yeast. *HOG1* is the yeast homolog of p38/MAPK14, a kinase that has been implicated in NFAT5 activation in some studies (though not in our experiments, **Figure S1D**).^28^ We disrupted both branches of the HOG pathway at three different levels using previously established strategies: disruption of *HOG1* itself, disruption of *PBS2*, encoding a MAPKK and scaffold protein, and the combined disruption of *SHO1*, *SSK2* and *SSK22* (**Figure 2J**).^13,35^ As expected, each of these mutations abrogated the activation of HOG pathway target genes in response to hypertonic stress (**Figure S2I**). However, in each of these three mutant backgrounds, mini-NFAT5 was still able to activate transcription of the 8TonE-*pCYC1*-GFP reporter in response to hypertonic stress, showing that the HOG pathway activity is dispensable for NFAT5 signaling in *S.cerevisiae* (**Figure 2K**).

Interestingly, the sensitivity of mini-NFAT5 to hypertonic stress was significantly enhanced in strains with disabled HOG signaling-- equivalent levels of 8TonE-*pCYC1*-GFP reporter activation were achieved at lower concentrations of added NaCI or sorbitol. Reporter activity *hoglΔ* cells declined at higher concentrations of NaCl, resulting in a bell-shaped dose response curve that qualitatively resembled the curve observed in mammalian cells (**Figure 1D**). The increased sensitivity of NFAT5 to hypertonic stress in *hog1Δ* cells may be caused by their inability to accumulate intracellular osmolytes to balance the additional extracellular NaCl. In addition, Hog1 also promotes Na^+^ efflux by phosphorylating the Na^+^/H^+^ antiporter Nha1.^36^ Thus, the loss of Hog1 compromises the ability of *S.cerevisiae* to maintain cytoplasmic ion homeostasis when confronted with increased extracellular ion concentrations. Consistent with this notion, the intracellular concentration of Na^+^ ions was elevated in *hog1Δ* cells, measured using either a fluorescent Na^+^ sensor or inductively coupled plasma optical emission spectroscopy (**Figures S2J** and **S2K**). Therefore the absence of a parallel osmoregulatory mechanism to adapt to hypertonic stress in *hog1Δ* cells results in a stronger stimulus for mini-NFAT5 activation.

### Intracellular ionic strength is the signal for NFAT5 activation

How can a nuclear protein like NFAT5 sense a change in the chemical potential of water in the extracellular space? Osmosensors in bacteria and fungi are often integral or peripheral membrane proteins.^12,37^ However, the specific physicochemical property sensed by osmosensors can vary between different organisms. The acute decrease in cell volume caused by hypertonic stress increases intracellular macromolecular crowding and consequently increases the phase separation propensity of many proteins (see **Figure 3A** for a chronology of changes triggered by hypertonic stress).^38–40^ Recent elegant work has shown that the WNK1 and ASK3 kinases, regulators of the RVI response that promote the intracellular accumulation of ions, and the plant transcriptional regulator SEUSS are sensors of macromolecular crowding that undergo phase separation in response to cell shrinkage (**Figure 3A**).^41–43^ Both cell shrinkage and RVI lead to an increase in intracellular ionic strength driven by an increase in the concentrations of ions (**Figure 3A**). Some bacterial osmosensors and osmoregulatory transporters are activated by the coulombic effect of an increase in the intracellular monovalent cation concentration caused by hypertonic stress.^44^ Sln1 and Sho1, membrane proteins in the HOG pathway, are thought to detect changes in turgor pressure or membrane tension.^31^ Each of these mechanisms-- macromolecular crowding, membrane tension, and ionic strength-- have been implicated in the activation of the mammalian NFAT5 pathway in different studies.^5,45–49^

**Figure 3.**
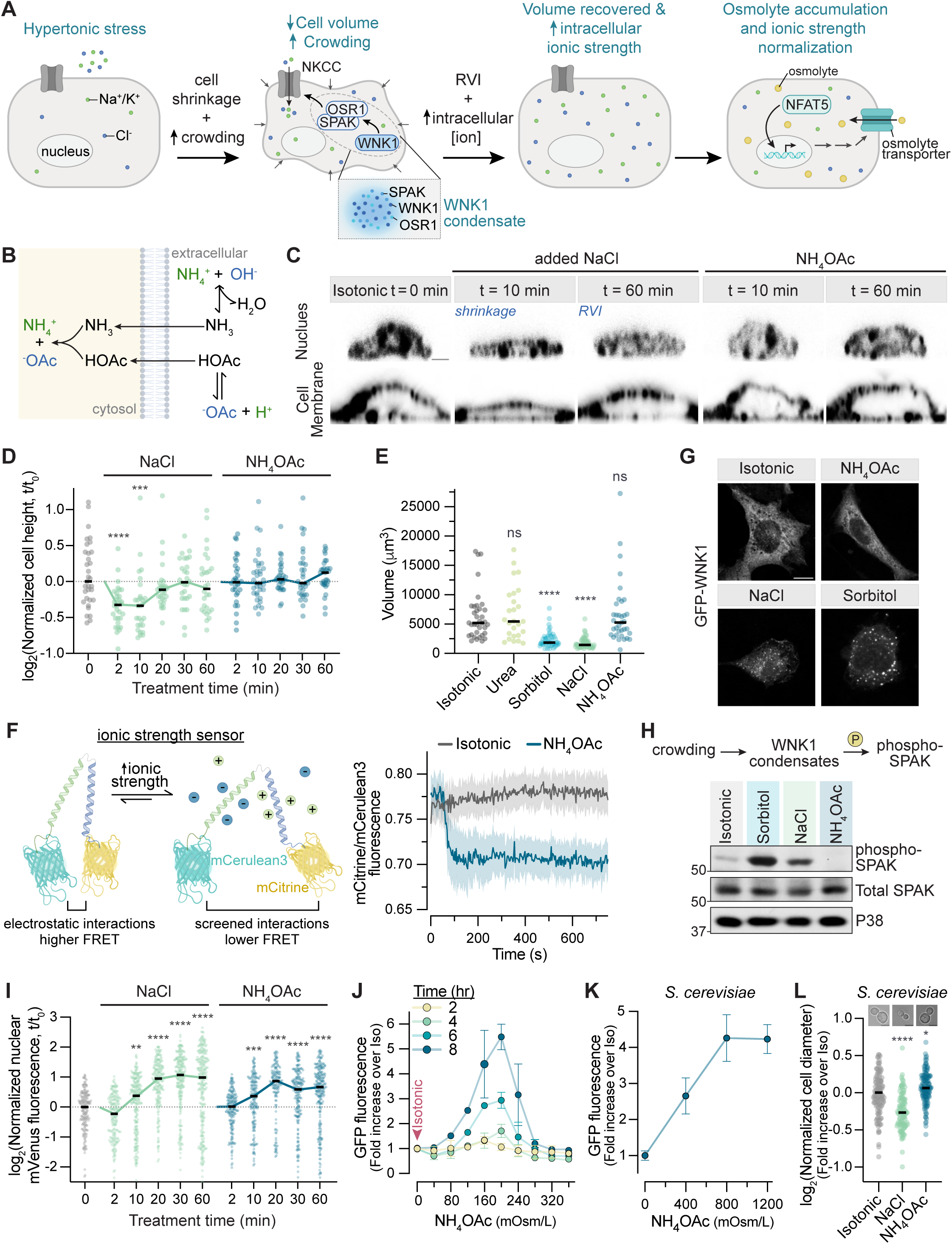
NFAT5 is activated by ionic stress. **(A)** Temporal sequence of cellular changes triggered by hypertonic stress. Hypertonic stress causes rapid cell shrinkage. The resulting increase in macromolecular crowding activates the regulatory volume increase (RVI) pathway: WNK1 initiates a kinase cascade that increases net influx of ions, leading to cell volume recovery within min, but at the cost of elevated intracellular ionic strength. If hypertonic stress persists, NFAT5 activation (over a slower timescale) allows the replacement of these excess ions with osmolytes, allowing restoration of the normal intracellular ion content. **(B)** Mechanism by which ammonium acetate (NH_4_OAc) crosses the plasma membrane and permeates cells. **(C)** Confocal images of IMCD3 cells treated with NaCl or NH_4_OAc (+200 mOsm/L added to isotonic media) and stained with DAPI to show nuclei (top) and CellMask to show the plasma membranes (bottom). Cells were imaged in the xz plane to show changes in cell height at 10 and 60 min after stress initiation.^91^ Scale bar: 2 μm. **(D)** The height of IMCD3 cells (*n*>28 per condition) was measured using confocal images of the type shown in **C** at various time points after addition of NaCl or NH_4_OAc (+200 mOsm/L) to isotonic media. **(E)** Volume of single IMCD3 cells (*n*>26 cells per condition) was measured using high-speed confocal imaging to capture z-stacks 10 min after the addition of 200 mOsm/L of urea, sorbitol, NaCl, or NH_4_OAc to isotonic media. **(F)** Cartoon showing the genetically-encoded Fluorescence Resonance Energy Transfer (FRET) sensor used to detect changes in intracellular ionic strength (left). Charged molecules like ions screen the attraction between the positively and negatively charged helices, reducing the FRET signal. Graph (right) shows the change in the mean mCitrine/mCerulean3 fluorescence ratio, along with an error envelope showing the SEM, from 20 individual IMCD3 cells after the addition of 200 mOsm/L NH_4_OAc to isotonic media. **(G)** Distribution of GFP fluorescence in *Wnk1*^-/-^ IMCD3 cells stably expressing GFP-WNK1 30 min after the addition of NH_4_OAc, NaCl, or sorbitol (+400 mOsm/L each) to isotonic media. **(H)** Immunoblotting was used to measure the abundance of phosphorylated SPAK and total SPAK in IMCD3 cells 30 min after the addition of NH_4_OAc, NaCl, or sorbitol (+400 mOsm/L each) to isotonic media. **(I)** Nuclear mVenus-NFAT5 fluorescence (*n*>145 IMCD3 cells per condition) at the indicated time points after the addition of NaCl or NH_4_OAc (+200 mOsm/L) to isotonic media. Representative images are shown in **Figure S3A**. These cells stably express full-length mVenus-NFAT5 (see **Figure 2B**). **(J)** 8TonE-GFP reporter activity in IMCD3-G cells was measured by flow cytometry at the indicated time points in response to increasing amounts of NH_4_OAc added to isotonic media. Each point shows the mean ± SD of three individual median measurements, each from a population of >2000 cells. **(K)** 8TonE-*pCYC1*-GFP reporter activity in yeast cells expressing mini-NFAT5 (**Figure 2D**) was measured by flow cytometry in response to increasing amounts of NH_4_OAc added to media (CSM). Each point shows the mean ± SD of six individual median measurements, each from a population of >5000 cells. **(L)** The diameter of yeast cells (*n*>98 per condition) was measured by microscopy 5 min after the addition of 1200 mOsm/L NaCl or NH_4_OAc to CSM. **Statistics**: Circles in (**D**,**E**,**I**,**L**) denote measurements from single cells and the black horizontal line marks the median of the population. The statistical significance of differences in comparison to the isotonic condition was determined by a Kruskal-Wallis test with Dunn’s multiple comparison test. P-value symbols are: **** *p*- value<0.0001, *** *p*-value<0.001, ** *p*-value<0.01, and * *p*-value<0.05. In (**D**), columns without symbols have *p*-values that are not significant. **See also Figure S3**.

**Supplementary Figure 3.**
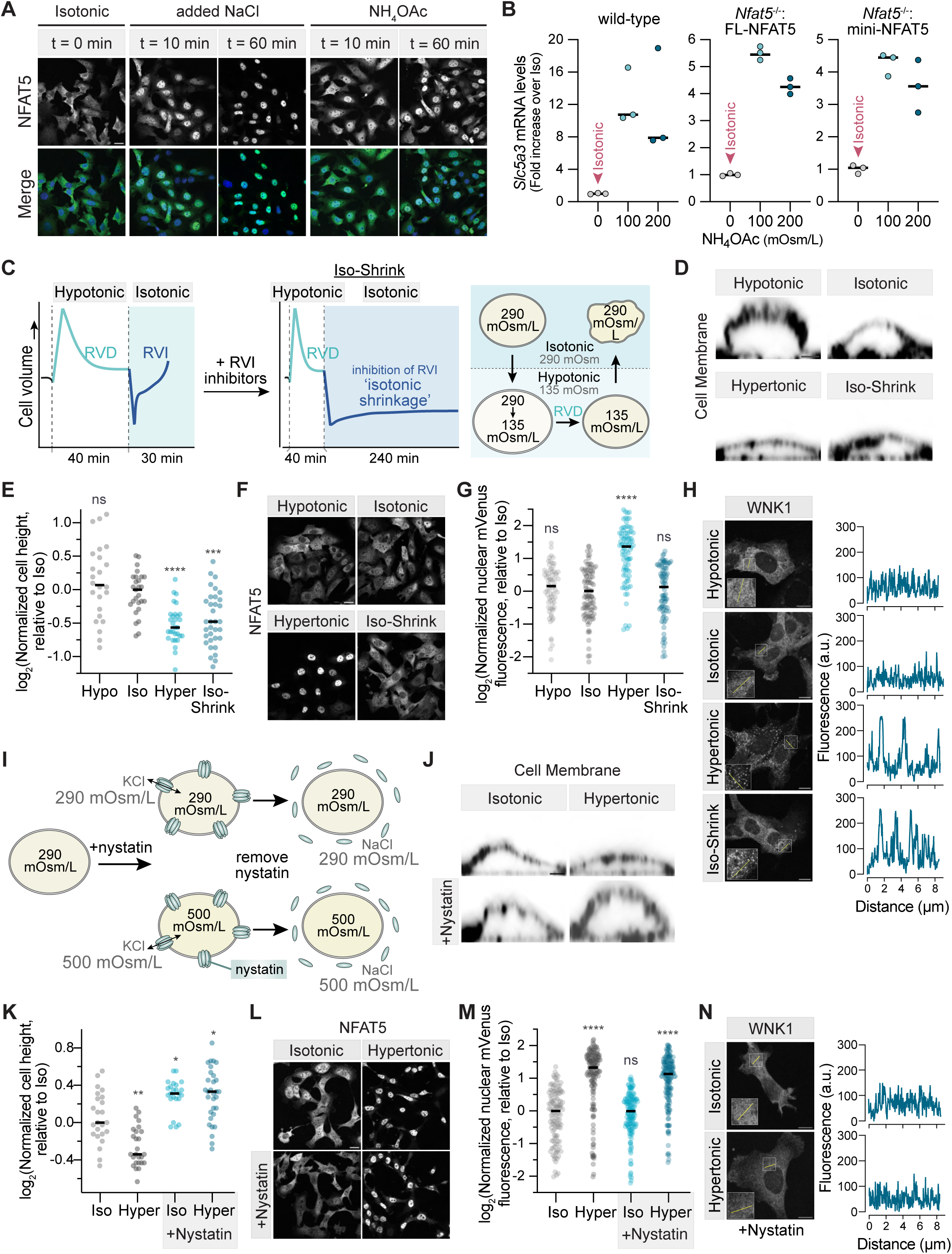
The effects of cell shrinkage and ionic stress on NFAT5 and WNK1, Related to Figure 3. **(A)** The subcellular localization of stably expressed mVenus-NFAT5 (top) after 10 or 60 min in hypertonic media (+200 mOsm/L NaCl or NH_4_OAc). Fluorescence signal from the mVenus tag fused to NFAT5 is shown alone (top) or merged with a nuclear stain (DAPI, bottom). Images like these were used for the quantitative analysis of NFAT5 nuclear accumulation shown in **Figure 3I**. Scale bar: 20 μm. **(B)** Expression of an NFAT5 target gene in WT IMCD3 cells or *Nfat5^-/-^* IMCD3 cells stably expressing mVenus-tagged full-length or mini-NFAT5 6 hrs after the addition of NH_4_OAc to isotonic media (see **Figure 3J** for a similar measurement using the 8TonE-GFP NFAT5 reporter). Black horizontal lines mark the median of three independent measurements, each shown as a point. **(C)** Diagram depicting the principle behind the isotonic shrinkage (Iso-Shrink) protocol, which allows cell shrinkage under isotonic conditions. Cells subjected to a hypotonic shock initially swell, but then restore their volume through regulatory volume decrease (RVD), which is caused by loss of osmotically active cytoplasmic ions and small molecules. The subsequent transfer of cells to isotonic media causes cell shrinkage since the cytoplasmic osmolarity is lower than media osmolarity. The addition of regulatory volume increase (RVI) inhibitors prevents cell volume recovery following isotonic shrinkage. Schematic taken from reference ^45^. **(D)** Confocal images of IMCD3 cells in the xz plane stained with CellMask to highlight the plasma membrane. Cells were exposed to hypotonic (∼200 mOsm/L), isotonic (∼300 mOsm/L), and hypertonic (∼500 mOsm/L with the addition of 200 mOsm/L NaCl) conditions, or subjected to the isotonic shrinkage protocol (bottom right Iso­Shrink panel) shown in **Figure S3C** and described in the methods. Scale bar: 2 μm. **(E)** The height of IMCD3 cells (*n*>24 cells per condition, measured from images of the type shown in **Figure S3D**) after the isotonic shrinkage protocol (**Figure S3C)** compared to cells in hypotonic, isotonic, or hypertonic media. **(F)** Subcellular localization of mVenus-NFAT5 stably expressed in IMCD3 cells following isotonic shrinkage, compared to cells in hypotonic, isotonic, or hypertonic media. Scale bar: 20 μm. **(G)** Nuclear mVenus fluorescence was measured (using images of the type shown in **Figure S3F**) from single IMCD3 cells (*n*>69 cells per condition) stably expressing full-length mVenus-NFAT5 following isotonic shrinkage, compared to cells in hypotonic, isotonic, or hypertonic media. **(H)** Distribution of GFP fluorescence in *Wnk1*^-/-^ IMCD3 cells stably expressing GFP-WNK1 after isotonic shrinkage, compared to its distribution in cells exposed to hypotonic, isotonic, or hypertonic media. Line scans show fluorescence intensity traces along the trajectories of the lines shown in the insets. Scale bar: 10 μm. **(I)** The ionophore nystatin forms ion conducting pores in the plasma membrane that allow equilibration of intracellular and extracellular osmolarity. When nystatin-treated cells are exposed to hypertonic media, intracellular ion concentrations rise <u>without</u> collateral cell shrinkage and macromolecular crowding. **(J)** Confocal images of IMCD3 cells in the xz plane stained with CellMask to highlight the plasma membrane. Cells were exposed to isotonic or hypertonic media in the presence or absence of nystatin. Scale bar: 2 μm. **(K)** The height of IMCD3 cells (*n*>23 cells per condition, measured from images of the type shown in **Figure S3I**) after exposure to isotonic or hypertonic media in the presence or absence of nystatin. **(L)** Subcellular localization of mVenus-NFAT5 in IMCD3 cells after exposure to isotonic or hypertonic media in the presence or absence of nystatin. Scale bar: 20 μm. **(M)** Nuclear mVenus fluorescence was measured (using images of the type shown in **Figure S3L**) from single IMCD3 cells (*n*>121 cells per condition) stably expressing full-length mVenus-NFAT5 after exposure to isotonic or hypertonic media in the presence or absence of nystatin. **(N)** Subcellular distribution of GFP-WNK1 following nystatin treatment in isotonic or hypertonic media. Line scans correspond to fluorescence intensity traces along the trajectories of the lines in the inset. Scale bar: 10 μm. **Statistics**: Circles in (**E**,**G**,**K**,**M**) denote measurements from single cells and the black horizontal lines mark the median of the populations. Statistical significance (**E**,**G**,**K**,**M**) of differences in comparison to the isotonic condition was determined by a Kruskal-Wallis testwith Dunn’s multiple comparison test (*n*>3 independent experiments). P-value symbols are: **** *p*-value<0.0001, *** *p*-value<0.001, ** *p*-value<0.01, and * *p*- value<0.05.

To identify the physicochemical signal that activates NFAT5, we sought to disentangle cell volume changes from changes in intracellular ionic strength. Addition of extracellular NaCl or sorbitol causes both cell shrinkage and increased ionic strength and so this treatment cannot be used to distinguish between these two signals. Nearly a century ago, cell physiologists made the observation that ammonium salts of weak acids like acetate and benzoate readily penetrate cells without causing a change in cell volume^50^ or intracellular pH^51^. When ammonium acetate (NH_4_OAc) is added to the extracellular medium, the uncharged products of its hydrolysis (NH_3_ and HOAc) enter cells and recombine to form the salt (**Figure 3B**). Thus, cell-permeant salts can increase intracellular ionic strength without causing cell shrinkage and its consequences of macromolecular crowding or changes in membrane tension. NH_4_OAc has been used to disrupt RNA condensation in cells without changing cell volume or intracellular pH.^52^

Unlike NaCl or sorbitol, NH_4_OAc did not cause shrinkage of IMCD3 cells but did increase intracellular ionic strength as measured by a genetically-encoded, FRET-based ionic strength sensor (**Figures 3C-3F**).^53^ NH_4_OAc also failed to induce either the phase separation or activation of the crowding sensor WNK1 (**Figures 3G** and **3H**).^41,54^ Thus, both morphometric and biochemical analyses confirmed that NH_4_OAc does not cause cell shrinkage or increase macromolecular crowding. In contrast, time-resolved experiments showed that NH_4_OAc, when added to cell culture media to increase osmolarity, activated NFAT5, as measured by nuclear accumulation and target gene transcription (**Figures 3I**, **3J**, **S3A**, and **S3B**). NH_4_OAc also triggered the activation of mini-NFAT5 in *S.cerevisiae*, despite not causing any changes in yeast cell size (**Figures 3K** and **3L**).

To validate our results with NH_4_OAc, we distinguished the effects of intracellular ionic strength from those of cell volume changes using two orthogonal, established methods. First, we used a protocol designed to isotonically shrink cells by cycling them from hypotonic to isotonic media in the presence of RVI inhibitors (**Figure S3C**).^41,45^ Isotonic shrinkage triggered WNK1 phase separation, showing that it increases macromolecular crowding, but failed to result in NFAT5 activation (**Figures S3D**-**S3H**). Second, we used nystatin, a reversible pore-forming ionophore, to increase the intracellular ion concentration without reducing cell volume.^55,56^ Conversely, under these conditions we observed NFAT5 activation, but no WNK1 phase separation (**Figures S3I-S3N**).

Hereafter we use the term “ionic stress” to describe the effect of NH_4_OAc on cells: increased intracellular ionic strength without an increase in the osmotic pressure gradient across the plasma membrane. While ionic stress can be a secondary consequence of hypertonic stress, the two are caused by distinct chemical changes in intracellular ionic strength and macromolecular crowding, respectively, and can be dissociated using specific experimental manipulations. Using three independent methods, our results established that NFAT5 activation is triggered by ionic stress. Cell-permeant salts like NH_4_OAc provide a simple way to increase intracellular ionic strength and impose ionic stress without altering cell volume and macromolecular crowding.

### NFAT5 forms biomolecular condensates in response to hypertonic and ionic stress

Analysis of the human NFAT5 sequence suggests that nearly the entire 1531 a.a. protein (with the exception of the 291 a.a. DBD) is intrinsically disordered (**Figures 4A** and **S4A**). The four other NFAT family proteins (NFAT1-4), all significantly shorter than NFAT5, are also predicted to be mostly disordered outside their DNA binding domains (**Figure S4B**). However, NFAT5 is unique amongst the NFAT proteins in that it possesses an unusually long predicted Prion-like Domain (PLD) embedded within its intrinsically disordered region (IDR) (**Figure 4A**). This ∼450 a.a. PLD is the fifth longest of the ∼240 PLDs found in the human genome.^57^ Deletion of the PLD in NFAT5 markedly diminishes its activity (**Figure S4C**). PLDs can function as sensors of the intracellular physicochemical environment, often switching between soluble and condensed or aggregated states in response to changes in pH, temperature or CO_2_.^58–61^

**Figure 4.**
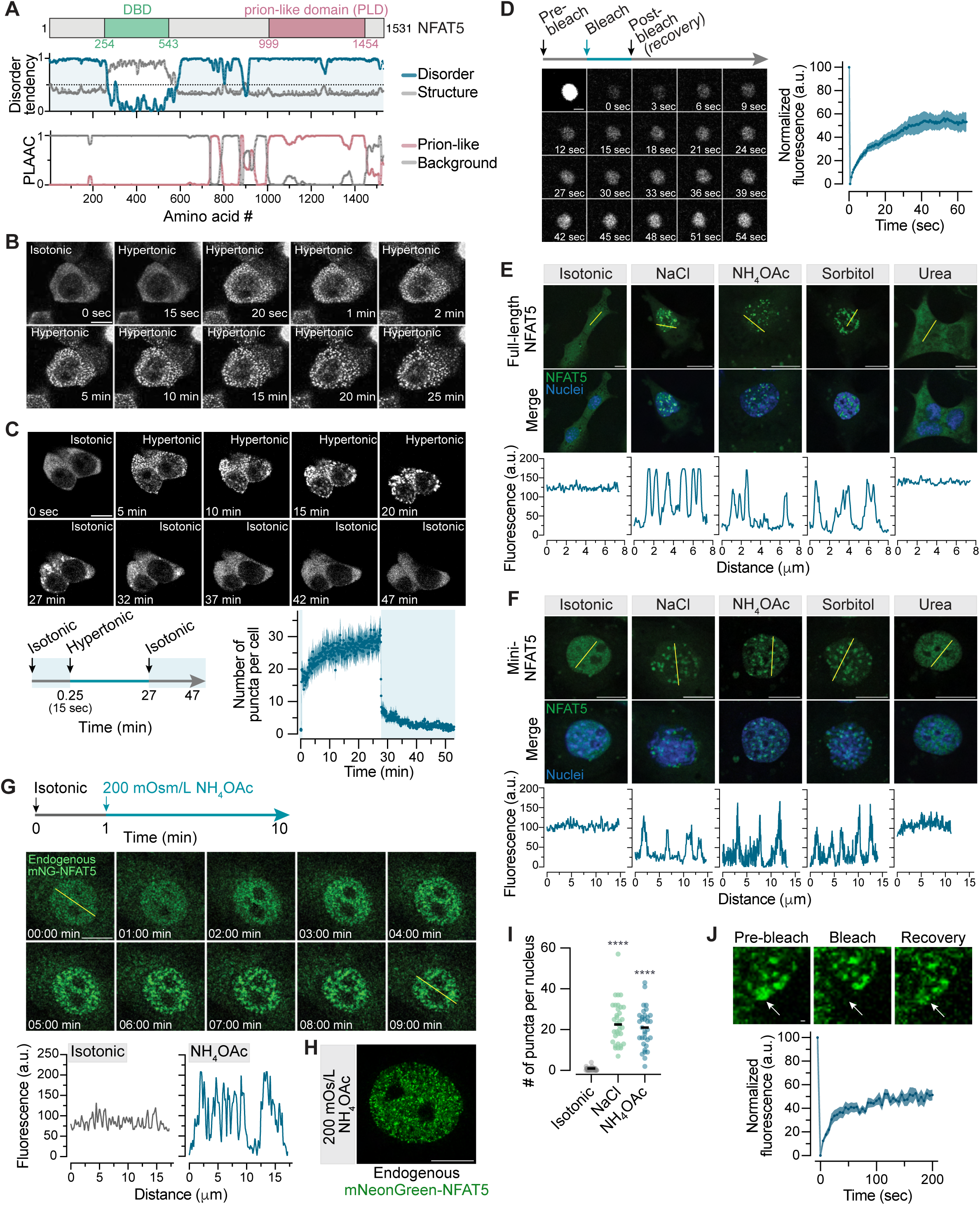
NFAT5 forms droplets in cells exposed to hypertonic or ionic stress. **(A)** Domain structure (top), predicted disorder tendency (middle) and predicted prion-like regions (bottom) of human NFAT5, all drawn on the same linear scale. NFAT5 has a predicted prion-like domain (PLD) and a structured DNA binding domain (DBD) embedded within intrinsically disordered regions (IDRs).^92,93^ **(B)** Snapshots from live cell imaging of 293T cells transiently transfected with GFP-NFAT5 and subjected to hypertonic stress. Isotonic media was replaced with hypertonic media (+100 mOsm/L NaCl) at t=15 seconds and images were collected for 25 min. **(C)** Live cell time course to test the reversibility of GFP-NFAT5 condensates in 293T cells. Cells were cycled from isotonic to hypertonic (+100 mOsm/L NaCl) and back to isotonic media according to the protocol shown on the bottom left. Mean (±SEM) number of droplets in 15 individual cells is shown on the graph (bottom right). Representative of three independent experiments. **(D)** Snapshots from a fluorescence recovery after photobleaching (FRAP) time course of a GFP-NFAT5 condensate in 293T subjected to hypertonic stress (+100 mOsm/L NaCl, 30 min). A corresponding recovery curve is shown on the right (mean ± SEM, *n*=9). **(E,F)** Subcellular distribution of full-length GFP-NFAT5 (**E**) or mVenus-mini-NFAT5 (**F**) stably expressed from a single locus in *Nfat5*^-/-^ IMCD3 cells (see methods) after the addition of NaCl, NH_4_OAc, sorbitol or urea (+200 mOsm/L each, 30 min) to isotonic media. The middle row of each panel shows NFAT5 distribution relative to nuclei. Line scans show fluorescence intensity traces along the trajectories of the yellow line in the images. **(G)** Live cell time course images of NH_4_OAc treated IMCD3 cells carrying NFAT5 tagged at its endogenous genomic locus with mNeonGreen (mNG) at the N-terminus in IMCD3 cells subjected to NH_4_OAc (see **Figures S5C**, **S5D** and **S5E** for characterization of this cell line). Line scans correspond to fluorescence intensity traces along the trajectories of the yellow lines shown in the images. **(H)** Maximum intensity projections of a live cell, super-resolution image (Structured Illumination Microscopy) of an IMCD3 nucleus carrying the endogenously tagged mNG-NFAT5. Cells were subjected to ionic stress (+200 mOsm/L NH_4_OAc) for 30 min prior to imaging. **(I)** Number of nuclear puncta in IMCD3 cells (*n*∼34 cells per condition) carrying endogenously-tagged mNG-NFAT5 after the addition of NaCl or NH_4_OAc (+200 mOsm/L, 30 min). Black horizontal line shows the median of each population and statistical significance of differences relative to the isotonic condition was determined using a Kruskal-Wallis testwith Dunn’s multiple comparison post-test (*n*>3 independent experiments). P-value symbol: **** *p*-value<0.0001. **(J)** FRAP recovery images and corresponding recovery curve (*n*=13; mean ± SEM) of endogenous mNG-NFAT5 puncta in IMCD3 cells subjected to ionic stress (+200 mOsm/L NH_4_OAc). Scale bars for panels (**B**,**C**,**E**,**F**,**G**,**H**): 10 μm; panels (**D**,**J**): 1 μm. **See also Figure S4 and S5**.

**Supplementary Figure 4.**
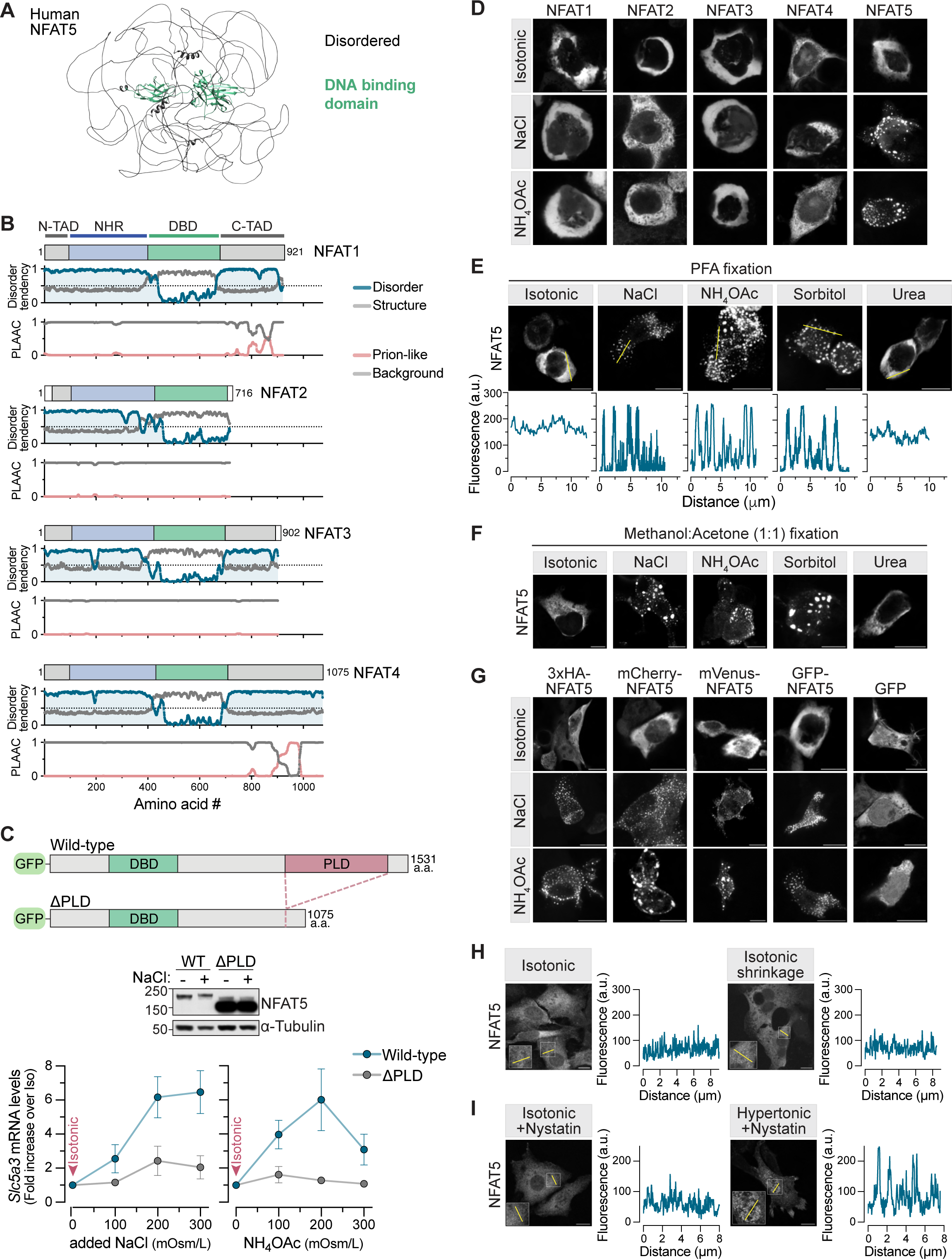
Formation of NFAT5 droplets in response to stress, Related to Figure 4. (**A**) AlphaFold prediction of the mostly disordered human NFAT5 (Uniprot ID: O94916) structure.^94^ (**B**) Domain structures, disorder tendency and predicted prion-like domains (PLDs) in human NFATs1-4 (compare to NFAT5 shown in **Figure 4A**). NFATs1-4 share a conserved NFAT-homology region (NHR) and DNA binding domain (DBD, also conserved with NFAT5) flanked by N-and C-terminal transactivation domains (TAD). (**C**) Expression of the NFAT5 target gene *Slc5a3* (bottom graphs) in *Nfat5*^-/-^ IMCD3 cells stably expressing wild-type GFP-NFAT5 (WT) or a variant lacking the PLD (GFP-NFAT5 APLD) 6 hrs after the addition of increasing amounts of NaCl (left) or NH_4_OAc (right) to isotonic media. Each point represents the mean ± SD of three independent measurements. Cartoon (top) shows the domain structure of the NFAT5 variants and immunoblot (middle) shows the abundances of each protein. (**D**) Distribution of GFP-tagged NFATs1-5 transiently transfected into 293T cells 30 min after the addition of NaCl or NH_4_OAc (+100 mOsm/L each) to isotonic media. (**E,F**) Distribution of GFP-NFAT5 in transiently-transfected 293T cells 30 min after the addition of NaCl, NH_4_OAc, sorbitol or urea (+100 mOsm/L each) to isotonic media. Cells were fixed with paraformaldehyde (PFA) in (**E**) or Methanol/Acetone in (**F**). Line scans in (**E**) correspond to fluorescence intensity traces along the trajectories of the yellow lines on the images. (**G**) Distribution of NFAT5 fused to different epitope tags in 293T cells after the addition of NaCl or NH_4_OAc (+100 mOsm/L each, 30 min) to isotonic media. Localization of GFP alone (without fusion to NFAT5) is shown in the rightmost column. (**H,I**) Distribution of GFP-NFAT5 stably expressed in *Nfat5*^-/-^ IMCD3 after isotonic shrinkage (**H**) or nystatin treatment (**I**) in the presence of an isotonic or hypertonic solution (see **Figures S3C** and **S3I** and Methods for details). Line scans correspond to fluorescence intensity traces along the trajectories of the lines shown in the insets. Scale bars for panels (**D-I**): 10 μm.

**Supplementary Figure 5.**
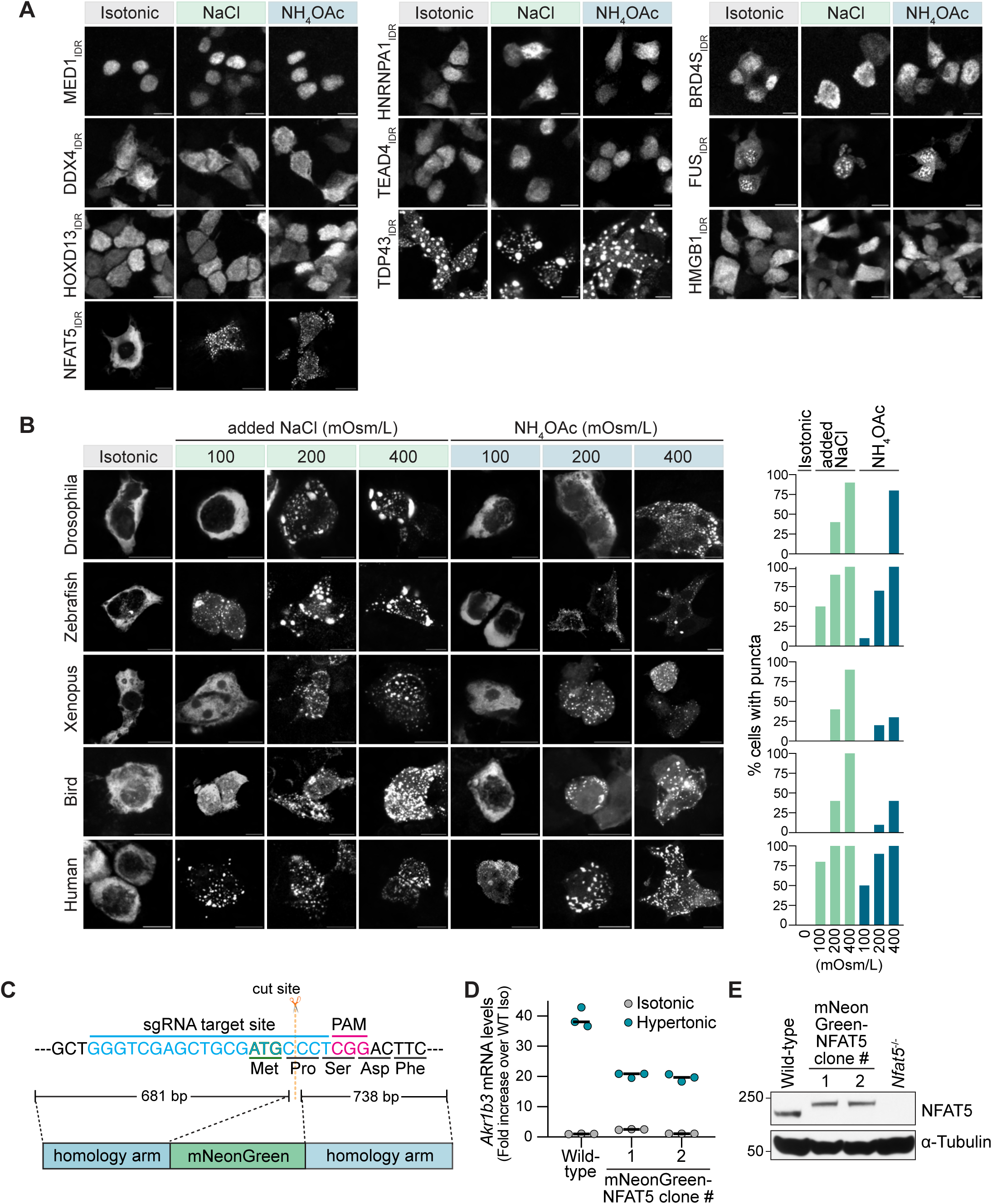
Evolutionarily conserved ionic stress sensing by NFAT5, Related to Figure 4. (**A**) Distribution of the GFP-tagged NFAT5 C-terminal IDR (CTD, **Figure 2A**) compared to Fluorescent Protein-tagged IDRs from nine unrelated proteins after transient transfection into 293T cells. Cells were fixed and imaged 30 min after the addition of NaCl or NH_4_OAc (+100 mOsm/L each) to isotonic media. (**B**) Distribution of chimeric NFAT5 proteins transiently transfected into 293T cells. Chimeras were generated by replacing the human NFAT5 CTD (a.a. 544-1531) with corresponding CTDs from insect (*Drosophila melanogaster*, a.a. 542-1210), fish (*Danio rerio*, a.a. 534-1257), amphibian (*Xenopus laevis*, a.a. 489-1408) and bird (*Columba livia*, a.a. 496-1455) NFAT5 homologs. Cells were imaged 30 min after addition of the indicated concentrations of NaCl or NH_4_OAc to isotonic media. Bar graphs on the right show the percentage of transfected cells with puncta at each salt concentration (*n*>210 cells evaluated per condition). (**C**) Exon 1 sequence of the mouse *Nfat5* gene targeted by the sgRNA used to insert the mNeonGreen coding sequence in IMCD3 cells using homology-directed repair. The Protospacer Adjacent Motif (PAM) and site of predicted cleavage by Cas9 are shown above the sequence; the donor template for homologous recombination is shown below. (**D**) Expression of the NFAT5 target gene *Akr1b3* in wild-type IMCD3 cells or two clonal *mNG-Nfat5* knock-in cell lines after 8 hrs in isotonic or hypertonic media (+200 mOsm/L NaCl). Black horizontal lines denote the median from three independent measurements shown as points. (**E**) Insertion of the mNG tag at the N-terminus of both *Nfat5* alleles in IMCD3 cells was confirmed by immunoblotting. Scale bars for panels (**A**,**B**): 10 μm.

Proteins with IDRs or PLDs have the propensity to engage in weak multivalent interactions, resulting in the formation of small networks or in segregative density transitions like phase separation or aggregation.^62^ IDRs are a prominent feature of most transcription factors and mediate transcriptional activation by enabling homo-and heterotypic multivalent interactions that mediate the formation of hubs or condensates.^63^ These associative behaviors, recently described by the unifying concept of “phase separation coupled to percolation” (PSCP), play regulatory roles in many different areas of cell biology.^62,64–66^ Given its sequence properties, we wondered whether NFAT5 displayed PSCP-like behaviors when cells were subjected to hypertonic stress. All five GFP-tagged NFAT proteins were homogeneously distributed in the cytoplasm when transiently over­expressed in 293T cells (**Figure S4D**). However, only NFAT5 (but not NFAT1-4) formed droplet-like condensates in response to hypertonic or ionic stress, both conditions that trigger NFAT5 activation (**Figures S4D** and **S4E**). Droplet formation was not observed when cells were exposed to the cell-permeable solute urea (**Figure S4E**). Droplets were seen in both live and fixed cells (**Figures 4B, S4E**, and **S4F**) and were independent of the tag attached to NFAT5 (**Figure S4G**). Droplet formation was rapid (**Figure 4B**) and was reversed (**Figure 4C**, **Video S1**) when cells were returned to isotonic media. Fluorescence recovery after photobleaching (FRAP) revealed that these NFAT5 droplets were dynamic, partially recovering on the time scale of minutes after bleaching (**Figure 4D**).

Isotonic shrinkage of cells, which increases macromolecular crowding without changing ionic strength, failed to induce the formation of NFAT5 droplets (**Figure S4H**), just as it failed to induce NFAT5 activation (**Figure S3G**). Conversely, increasing intracellular ionic strength without changing cell volume by using nystatin (**Figure S4I**) or adding NH_4_OAc (**Figure S4E**) induced the formation of NFAT5 droplets. Thus, NFAT5 droplet­formation was driven by increased cytoplasmic ionic strength, rather than by an increase in macromolecular crowding.

We considered whether the response of NFAT5 to ionic stress was shared with other IDR-containing proteins since increased intracellular sodium has been shown to increase the liquidity of ASK3 condensates.^67^ Hypertonic and ionic stress did not alter the condensation behavior of IDRs from nine other transcriptional regulators that undergo PSCP in various contexts, suggesting that this is a property unique to NFAT5 (**Figure S5A**).^68,69^ Ionic stress sensitivity also seems to be an evolutionarily conserved property of the NFAT5 proteins, since it was retained when the CTD of human NFAT5 was replaced with that from bird, fish, amphibian and even invertebrate NFAT5 homologs (**Figure S5B**). Interestingly, IDRs from different species demonstrated varying sensitivity to hypertonic and ionic stress, perhaps reflecting the fact that NFAT5 is tuned to respond to the specific stress ranges faced by each organism.

The transient expression of NFAT5 in 293T cells provides a rapid, easy-to-score assay to evaluate how various treatments change its propensity to self-associate. However, NFAT5 is over-expressed under these conditions and not properly regulated since it does not accumulate in the nucleus in response to hypertonic stress.^17^ Consequently, we also tested whether ionic stress-induced NFAT5 condensation was observed in the nucleus at lower, more physiological protein abundances. First, we used *Nfat5*^-/-^ IMCD3 cells stably expressing GFP-tagged NFAT5 or mVenus-tagged mini-NFAT5 from a single genomic locus-- both of which can restore transcriptional responses to hypertonic stress to cells lacking endogenous NFAT5 (**Figure 2C**). NFAT5 accumulated in the nucleus properly in these cells and formed nuclear puncta in response to hypertonic or ionic stress (**Figure 4E**, **Video S2**). While mini-NFAT was designed to be constitutively nuclear (**Figure 2A**), this nuclear pool, which is diffusely distributed under isotonic conditions, formed puncta in response to hypertonic or ionic stress (**Figure 4F**). Thus, like its transcriptional activity (**Figure 2C**), the propensity of NFAT5 to self-associate in the nucleus can be influenced by extracellular hypertonicity.

To examine the behavior of endogenous NFAT5 expressed from its native promoter, we introduced an in-frame mNeonGreen (mNG) tag at the N-terminus of both *Nfat5* alleles in IMCD3 cells (**Figure S5C**). Live cell imaging revealed that endogenous mNG-NFAT5 accumulated in the nucleus, formed puncta and activated target genes in response to ionic and hypertonic stress (**Figures 4G-4I**, **S5D** and **S5E**, **Video S3**). Interestingly, NFAT5 nuclear puncta were polymorphic in shape (**Figure 4G** and **4J**), in contrast to the more spherical droplets seen with overexpressed, cytoplasmic NFAT5 in 293T cells. Despite this morphological difference, mNG-NFAT5 nuclear puncta were dynamic structures as assessed using FRAP (**Figure 4J**).

### An intrinsically disordered, prion-like region mediates ionic strength sensing by NFAT5

Like full-length NFAT5, the isolated CTD or PLD of NFAT5 both formed droplets in response to hypertonic or ionic stress (**Figures 5A** and **5B**). Could these regions of NFAT5 endow a heterologous protein with sensitivity to ionic stress?

**Figure 5.**
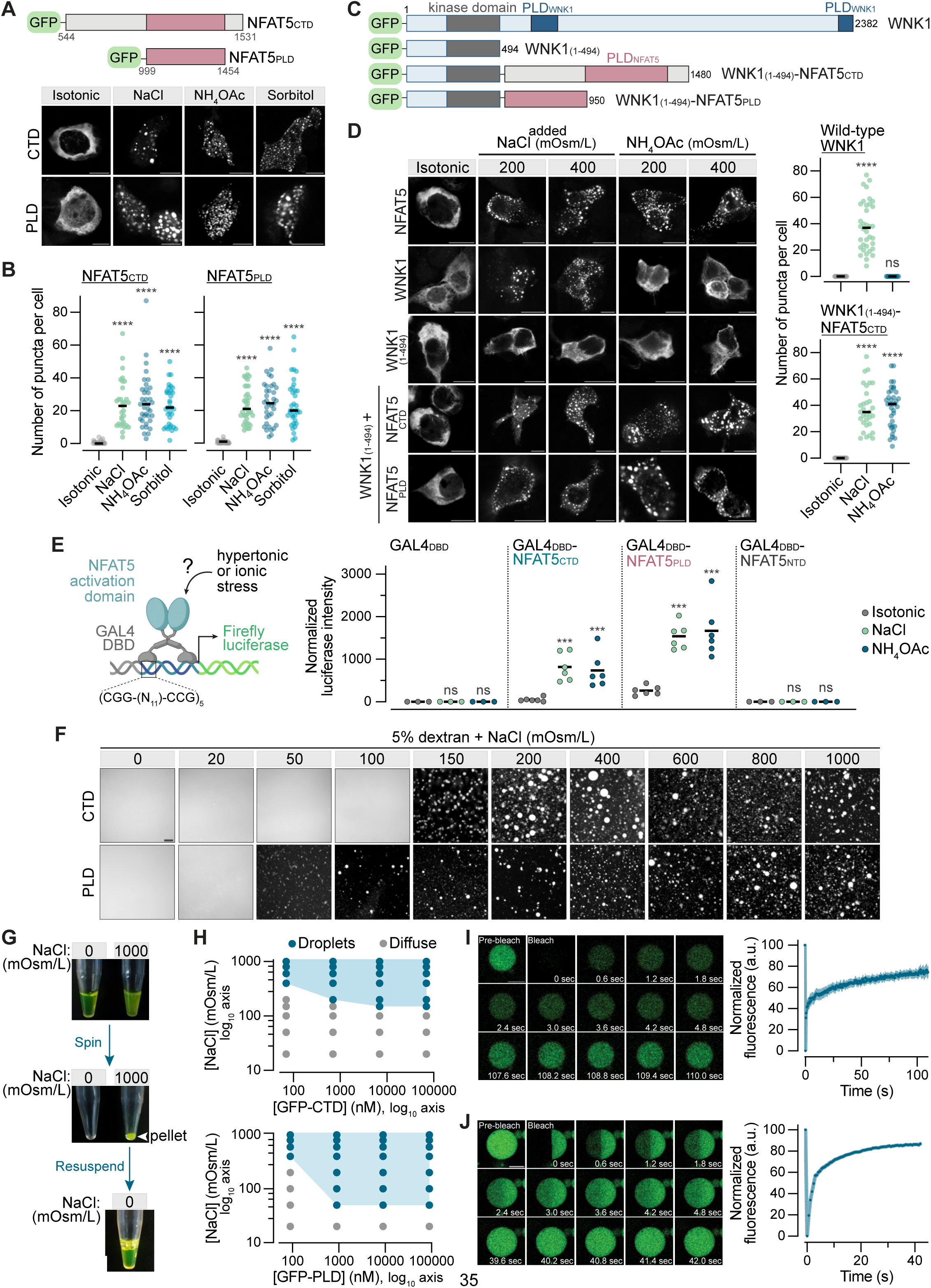
The C-terminal intrinsically disordered region of NFAT5 can sense solution ionic strength. (**A, B**) Distribution (**A**) of GFP-CTD (C-terminal domain of NFAT5, a.a. 544-1531) or GFP-PLD (prion-like domain of NFAT5, a.a. 999-1454) in 293T cells 30 min after the addition of NaCl, NH_4_OAc or sorbitol (+100 mOsm/L each) to isotonic media (see **Figure 2A** for the position of the CTD and PLD in full-length NFAT5). Images of the type shown in (**A**) were used to measure the number of puncta per cell (*n*>25 cells, with black horizontal line showing the median and each point showing the number of puncta in one cell). (**C**) Domain structure of WT and chimeric GFP-tagged WNK1 constructs. The WNK1 1-494 fragment contains a fully intact kinase domain and a C-terminal (a.a. 495-2382) IDR that senses macromolecular crowding. In the chimeras, the IDR of WNK1 is replaced with the CTD or the PLD of NFAT5. (**D**) Distribution of full-length and chimeric GFP-tagged WNK1 constructs (shown in **C**) in 293T cells 30 min after the addition of the indicated concentrations of NaCl or NH_4_OAc to isotonic media. The graphs (right) show the number of puncta in cells treated with NaCl or NH_4_OAc (200 mOsm/L, 30 min) (*n*>20 cells with black horizontal line showing the median and each point showing the number of puncta in one cell). (**E**) Design of synthetic transcription factors through fusion of the DNA binding domain (DBD, a.a. 1-147) of the *S. cerevisiae* GAL4 protein to either the NFAT5 CTD, PLD or N-terminal domain (NTD, **Figure 2A**) (left). Each of the GAL4 fusions were tested for their abilities to activate a firefly luciferase reporter gene in response to NaCl (+200 mOsm/L) or NH_4_OAc (+100 mOsm/L). Solid horizontal lines denote mean values calculated from >3 independent measurements shown as points. (**F**) Fluorescence microscopy was used to assess droplet formation *in vitro* by purified (**Figure S6A**) GFP-CTD (70 μM, top row) and GFP-PLD (90 μM, bottom row) in buffer containing 5% dextran and the indicated concentrations of NaCl. (**G**) Reversibility of GFP-CTD condensates assessed by a centrifugation and resuspension assay. Condensates (which cause turbidity, top panel) triggered by the addition of NaCl to a solution of 70 μM GFP-CTD can be sedimented (middle panel). This condensate pellet readily dissolves to a clear solution in a low ionic strength buffer (bottom panel). All solutions contain 5% dextran. (**H**) Phase diagrams for purified GFP-CTD (top) and GFP-PLD (bottom). Blue shading denotes conditions where the GFP fluorescence was exclusively in droplets while the unshaded area encompasses conditions where GFP fluorescence remained diffuse (see **F** for examples of each). The boundary between the shaded and unshaded areas of the graph is taken as the phase boundary; crossing this boundary leads to the abrupt drop of diffuse fluorescence and emergence of droplets. Images were obtained across three replicates per condition. (**I**,**J**) Full-bleach (**I**) and half-bleach (**J**) FRAP recovery images and corresponding recovery curves (*n*=10; mean ± SEM) of GFP-CTD condensates assembled in buffer with 5% dextran, 200 mOsm/L NaCl and 70 μM protein. Scale bars for panels (**A**,**D**): 10 μm; panels (**F**,**I**,**J**): 5 μm. **Statistics**: Statistical significance of the differences in comparison to the isotonic condition was determined in (**B,D**) with a Kruskal-Wallis test and Dunn’s multiple comparison test and in (**E**) with a two-way ANOVA test and Sidak’s multiple comparison test (*n*>3 independent experiments). P-value symbols are: **** *p*- value<0.0001 and *** *p*-value<0.001. **See also Figure S6**.

**Supplementary Figure 6.**
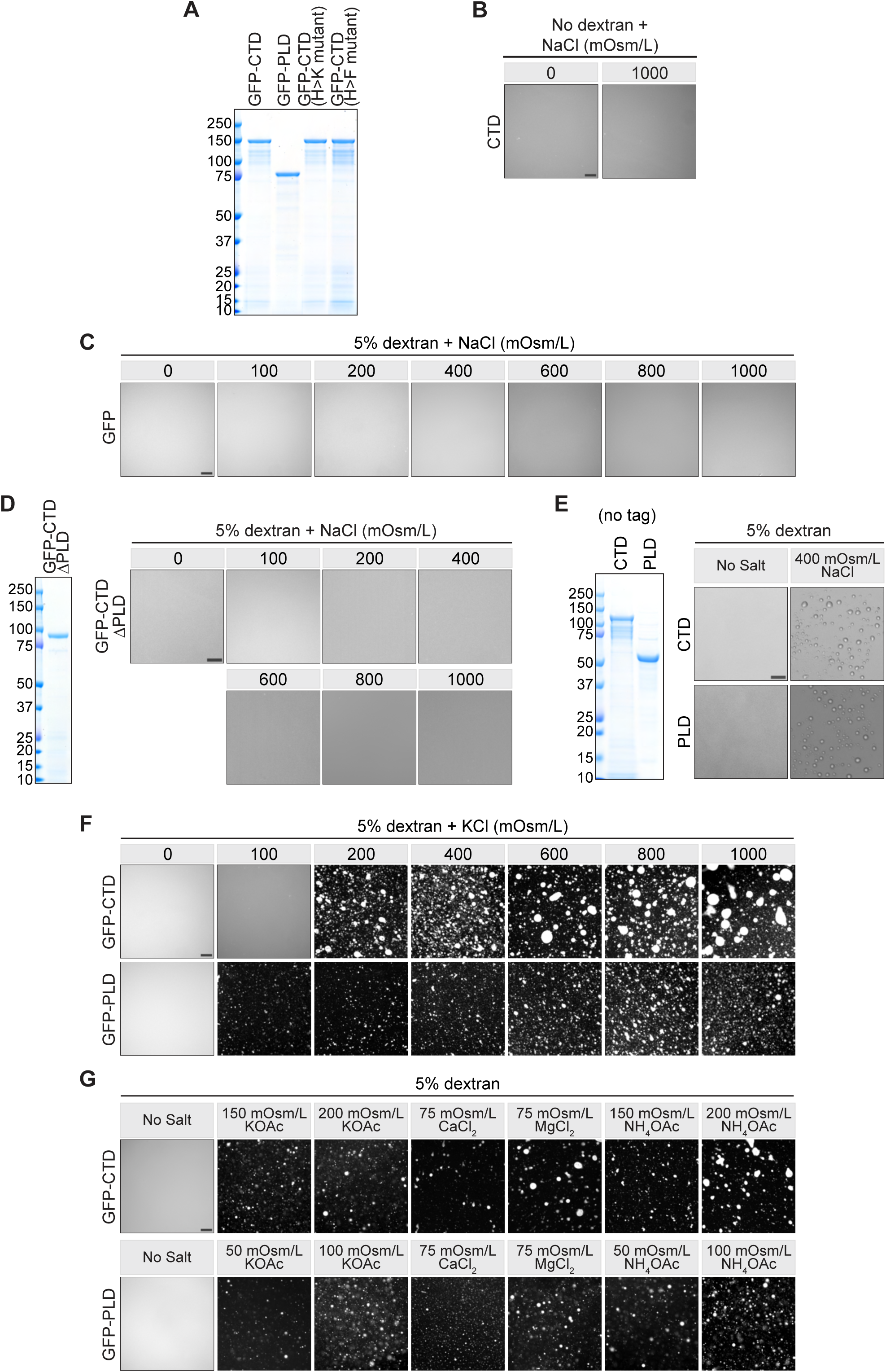
Characterization of the *in vitro* droplet formation assay of NFAT5 protein fragments, Related to Figure 5. **(A,D,E)** Coomassie-stained polyacrylamide gels showing the purity of proteins used for *in vitro* droplet formation assays in **Figures 5**, **6** and **7**. Molecular weight standards in kilodaltons (kDa) are indicated to the left. (**B**) Fluorescence microscopy was used to assess droplet formation *in vitro* by purified GFP-CTD (70 μM) in buffered solution <u>without</u> Dextran. (**C,D**) Fluorescence microscopy was used to assess droplet formation *in vitro* by 100 μM purified GFP or GFP-CTD-APLD (CTD lacking the PLD, see **Figure S4C**) at the indicated concentrations of NaCl. (**E**) Brightfield microscopy was used to assess droplet formation by untagged NFAT5 CTD (70 μM) and PLD (90 μM). (**F,G**) *In vitro* droplet formation by GFP-NFAT5 CTD (70 μM, top row) or GFP-NFAT5 PLD (90 μM, bottom row) at increasing concentrations of KCl (**F**) or in the presence of a variety of different salts (**G**). All solutions contained 5% dextran. Scale bars for panels (**B-G**): 5 μm.

As a first test case, we replaced the C-terminal domain of WNK1, previously identified as a sensor of macromolecular crowding in the cytoplasm, with either the CTD or the PLD of NFAT5 (**Figure 5C**).^41^ WNK1 phase separates in response to cell shrinkage and increased macromolecular crowding, induced by hypertonic stress, but not in response to ionic stress (**Figures 5C** and **5D**). In contrast, NFAT5 droplets form in response to elevated ionic strength, but not cell shrinkage. Chimeric WNK1-NFAT5 proteins formed droplets in response to both hypertonic and ionic stress, showing that transplantation of the NFAT5 CTD or PLD could transfer ionic strength sensing to WNK1 (**Figures 5C** and **5D**). The WNK1-NFAT5 chimeras did not activate the kinase cascade downstream of WNK1, perhaps because NFAT5 condensates do not provide the appropriate environment for the recruitment or activation of downstream WNK1 pathway components.

Second, inspired by previous studies, we constructed synthetic transcription factors by fusing the DBD of the *S. cerevisiae* GAL4 protein to either the NFAT5 CTD or PLD (**Figure 5E**).^32,70^ The activities of these GAL4_DBD_ fusions were tested using a reporter construct containing 5 binding sites for GAL4 positioned upstream of a minimal promoter driving the firefly luciferase gene. The GAL4_DBD_ alone failed to activate luciferase expression. However, both the GAL4_DBD_^_^NFAT5_CTD_ and GAL4_DBD_^_^NFAT5_PLD_ chimeras activated the luciferase reporter in response to ionic stress. These experiments show that the NFAT5 CTD and PLD are sufficient to activate transcription of target genes in response to increases in intracellular ionic strength. In addition, these results show that ionic stress likely controls a step in transcriptional activation different from DNA binding.

To directly demonstrate that NFAT5 can sense ionic strength without other cellular factors, we expressed GFP-fusions of its CTD and PLD domains in *E.coli* and purified each protein in a low ionic strength buffered solution (20mM Na-HEPES) for *in vitro* phase separation assays (**Figure S6A**). In the absence of any added salt, 5% dextran, a polysaccharide commonly used to mimic the crowded milieu of the cytoplasm in such assays, failed to induce phase separation (**Figure 5F**). However, gradually elevating ionic strength abruptly triggered phase separation of both GFP-CTD and GFP-PLD across a narrow window (∼50 mOsm/L) of NaCl concentrations: the diffuse, homogeneous GFP fluorescence coalesced into spherical droplets, concomitantly reducing background fluorescence (**Figure 5F**). The sharp transition between diffuse and condensed GFP-CTD and GFP-PLD supports the crossing of a phase boundary rather than titration of an ion­binding site in the protein. The GFP-CTD condensates could be isolated by centrifugation and readily dissolved when resuspended in a low ionic strength buffer, indicating that they were formed by dynamic, reversible interactions and were not irreversible aggregates (**Figure 5G**).

Several controls were done to ensure the specificity of this phenomenon. First, phase separation required the presence of dextran as a crowding agent: no droplets were observed without dextran at NaCl concentrations as high as 1000 mOsm/L (**Figure S6B**). Second, GFP alone (**Figure S6C**) or a GFP-CTD protein lacking the PLD (**Figure S6D**) did not form droplets. Notably, excision of the PLD from NFAT5 also abrogates its transcriptional activity (**Figure S4C**). CTD and PLD proteins lacking the GFP tag also formed droplets as assessed by brightfield microscopy (**Figure S6E**). Finally, phase separation did not specifically require NaCl but could be triggered by increasing the ionic strength of the solution using a variety of different salts (**Figure S6F** and **S6G**).

Phase diagrams of GFP-CTD and GFP-PLD were constructed by varying the ionic strength at protein concentrations ranging between ∼100 nM and ∼100 μM (**Figure 5H**). Phase boundaries of both proteins were distinct: GFP-PLD was more sensitive to ionic strength at all protein concentrations tested. At the lowest protein concentrations tested (∼70-100 nM), phase separation was observed only at NaCl concentrations above ∼300 mOsm/L (**Figure 5H**). Thus, the purified NFAT5 CTD, when present at near-endogenous nanomolar concentrations *in vitro*, phase separates in response to NaCl concentrations that would impose hypertonic stress on cells.

We assessed the material properties of GFP-CTD condensates using FRAP. Full-bleach experiments showed that there was clearly exchange between the diffuse and condensed phases (**Figure 5I**). Kinetics of exchange were multi-phasic and recovery was incomplete, suggesting the presence of multiple populations of GFP-CTD in the condensates that exchanged with the diffuse phase at different rates. Half-FRAP experiments also demonstrated recovery of GFP-CTD fluorescence in the bleached half (**Figure 5J**). However, most of this recovery was due to exchange with the diffuse phase rather than due to internal mixing within the condensate, since there was no concomitant decline in the fluorescence of unbleached half.^71^

### NFAT5 condensation propensity correlates with its transcriptional activity

Ionic strength stimulates the transcriptional activity of NFAT5 in cells and also promotes the self­association of NFAT5 into condensates, both *in vitro* under purified conditions and in intact cells. These observations suggest the hypothesis that these two properties are linked-- ionic-strength triggered self­association (and perhaps association with other IDR-containing proteins) drives transcriptional activation. Such multivalent interactions (which may or may not result in a segregative transition like phase separation) have been recently proposed to explain the transactivation capacity of IDRs that are a universal feature of transcription factors.^63^ The unique feature of NFAT5 would be the presence of an IDR that can reversibly engage in such associative interactions in response to the ionic strength in the nucleoplasm.

We sought to perturb the chemical interactions that drive NFAT5 condensation using three methods with increasing degrees of specificity. The aliphatic alcohol 1,6-hexanediol disrupts the weak hydrophobic interactions that often drive associative interactions between IDRs.^72^ Treatment of cells with 1,6-hexanediol inhibited the formation of NFAT5 droplets (**Figure 6A**) and the transcription of NFAT5 target genes (**Figures 6B** and **S7A**). The use of aliphatic alcohols has been criticized because these nonspecific agents can potentially disrupt many cellular processes. Consequently, we turned to making mutations in the a.a. sequence of NFAT5 itself. Compared to the human proteome, the NFAT5 sequence is enriched in glutamine (Q) and serine (S), but depleted in the charged residues glutamic acid (E), aspartic acid (D), lysine (K) and arginine (R) (**Figures 6C** and **S7B**). Given the well-established importance of glutamine residues for the structural and functional properties of PLDs, we mutated all 109 glutamine residues in the NFAT5 PLD to alanine. This variant, hereafter called “NFAT5_QA”, was expressed at levels comparable to WT NFAT5 (**Figure S7C**), but failed to accumulate in the nucleus, form puncta (**Figures 6D** and **6E**), and activate target genes (**Figure 6F**) in response to hypertonic stress. In contrast, changing all 69 of the serine residues to alanine in the PLD had no effect on NFAT5 activity (**Figures S7E-S7I**).

**Figure 6.**
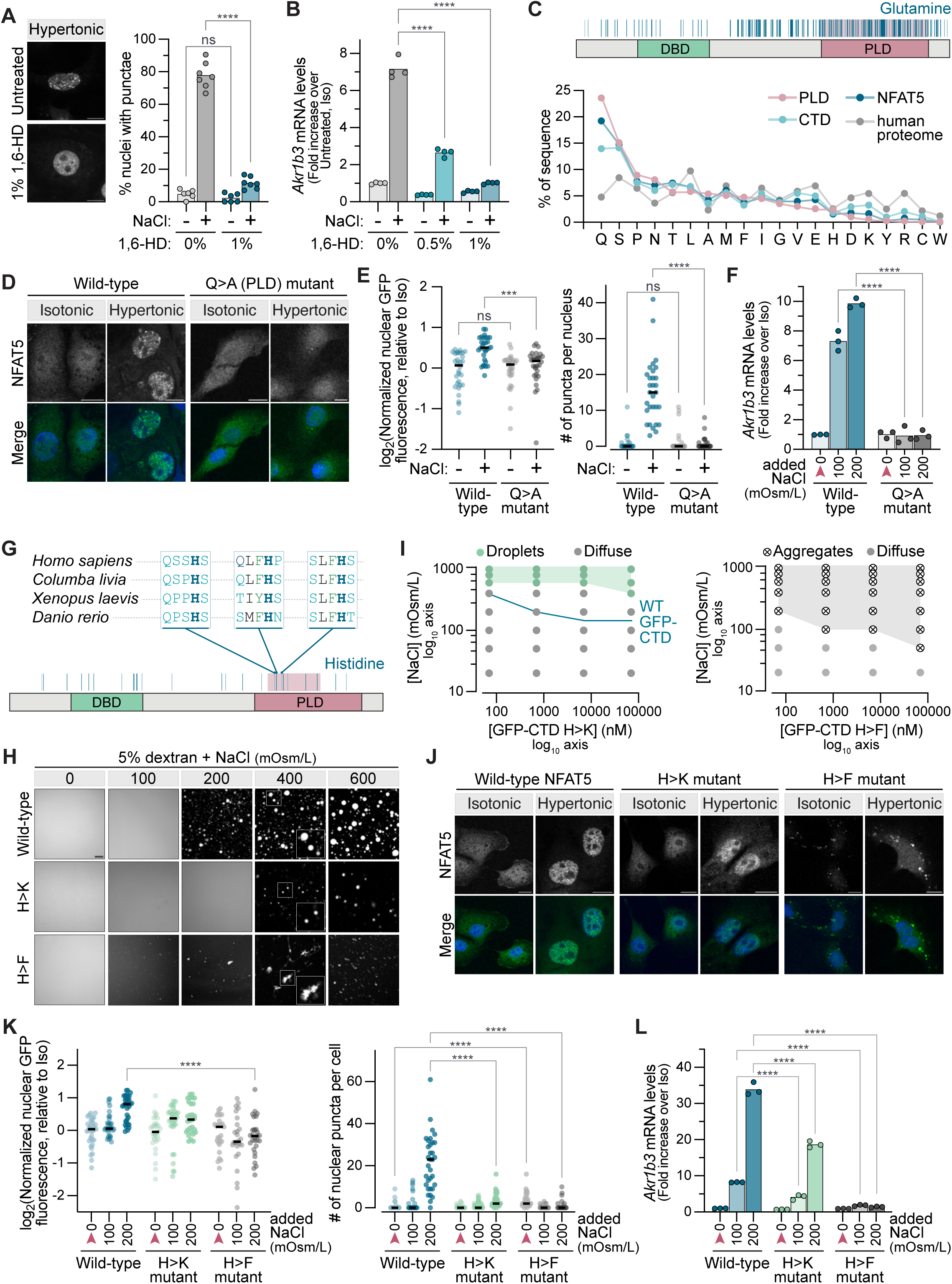
NFAT5 activity correlates with phase separation propensity. **(A)** Distribution of nuclear GFP-NFAT5 stably expressed in *Nfat5*^-/-^ IMCD3 cells exposed to hypertonic stress (+200 mOsm/L NaCl added to isotonic media) in the presence or absence of 1% 1,6-hexanediol (1,6-HD) (left). The percentage of cells with nuclear puncta was calculated from >100 cells per point shown; the bar shows the mean of 6-7 independent measurements (right). **(B)** Expression of the NFAT5 target gene *Akr1b3* in response to a 6 hr treatment of 0, 0.5, or 1% 1,6-HD in isotonic or hypertonic (+200 mOsm/L NaCl) media. Bars denotes the mean of 4 independent experiments shown as points. **(C)** Domain structure of NFAT5 with the positions of all glutamine residues marked with vertical blue lines (top). Amino acid composition in NFAT5 or its isolated CTD and PLD fragments compared to the composition in the human proteome (bottom). (**D,E**) Subcellular distribution (**D**) of full-length GFP-NFAT5 or the GFP-NFAT5_QA mutant (all 109 glutamine residues in the PLD mutated to alanine) stably expressed from a single locus in *Nfat5*^-/-^ IMCD3 cells after the addition of NaCl (+200 mOsm/L, 30 min) to isotonic media. Fluorescence signal from NFAT5 is shown alone (top) and merged with DAPI signal to show nuclei (bottom). Images of the type shown in (**D**) were used in (**E**) to measure nuclear NFAT5 fluorescence (left) and number of NFAT5 nuclear puncta (right) per cell (*n*>30 cells, with black horizontal line showing the median and each point showing the measurement in one cell). (**F**) Expression of the NFAT5 target gene *Akr1b3* following the addition of NaCl to isotonic media (red arrowhead) for 8 hrs. (**G**) Domain structure of NFAT5 with the positions of all histidine residues marked by vertical blue lines. Red shading highlights the seven histidines within the PLD targeted for mutagenesis. Cross-species sequence conservation of a subset of these histidines is shown (top). (**H**) *In vitro* droplet formation propensity of purified (**Figure S6A**) GFP-CTD (70 μM, top row), GFP-CTD_HK (70 μM) and GFP-CTD_HF (70 μM) at the indicated NaCl concentrations. The seven histidines in the NFAT5 PLD highlighted in (**G**) were mutated to lysines or phenylalanines in GFP-CTD_HK and GFP-CTD_HF, respectively. Magnified insets show droplet shape (uniformly spherical for GFP-CTD and GFP-CTD_HK and both irregular and polymorphic for GFP-CTD_HF) (**I**) Phase diagrams for purified GFP-CTD_HK (left) and GFP-CTD_HF (right) (see **Figure 5H** for description). Shaded areas mark conditions where GFP-CTD_HK (green shading, left) or GFP-CTD_HF (gray shading, right) form droplets or aggregates, respectively. The blue trace indicates the position of the boundary between diffuse and droplet phases of WT GFP-CTD (from **Figure 5H**). Images were obtained across 3 biological replicates per condition. (**J,K**) Subcellular distribution (**J**) of full-length GFP-NFAT5 or the corresponding HK and HF mutants (see **G**) stably expressed from a single locus in *Nfat5*^-/-^ IMCD3 cells after the addition of NaCl (+200 mOsm/L, 30 min) to isotonic media. Fluorescence signal from NFAT5 is shown alone (top) and merged with DAPI to mark nuclei (bottom). (**K**) Images of the type shown in (**J**) were used to measure the nuclear NFAT5 fluorescence (left) and number of NFAT5 nuclear puncta (right) per cell (*n*>30 cells, with black horizontal line showing the median and each point showing the measurement in one cell). Red arrowheads mark the isotonic condition. (**L**) Expression of the NFAT5 target gene *Akr1b3* 8 hrs after the addition of NaCl to isotonic media (red arrowheads). Bars denote the mean of 3 measurements, each shown as points. Representative of 4 independent experiments. In panels (**F**,**K**,**L**) red arrowheads indicate the isotonic (300 mOsm/L) condition. Scale bars for panels (**D**,**J**): 10 μm; panel (**H**): 5 μm. **Statistics**: For (**A**,**B**,**F**,**L**), statistical significance was determined by a two-way ANOVA test, Sidak’s multiple comparison. For (**E**,**K**) statistical significance was determined by a Kruskal-Wallis test, Dunn’s multiple comparison. P-value symbols are: **** *p*-value<0.0001 and *** *p*-value<0.001. **See also Figure S7**.

**Supplementary Figure 7.**
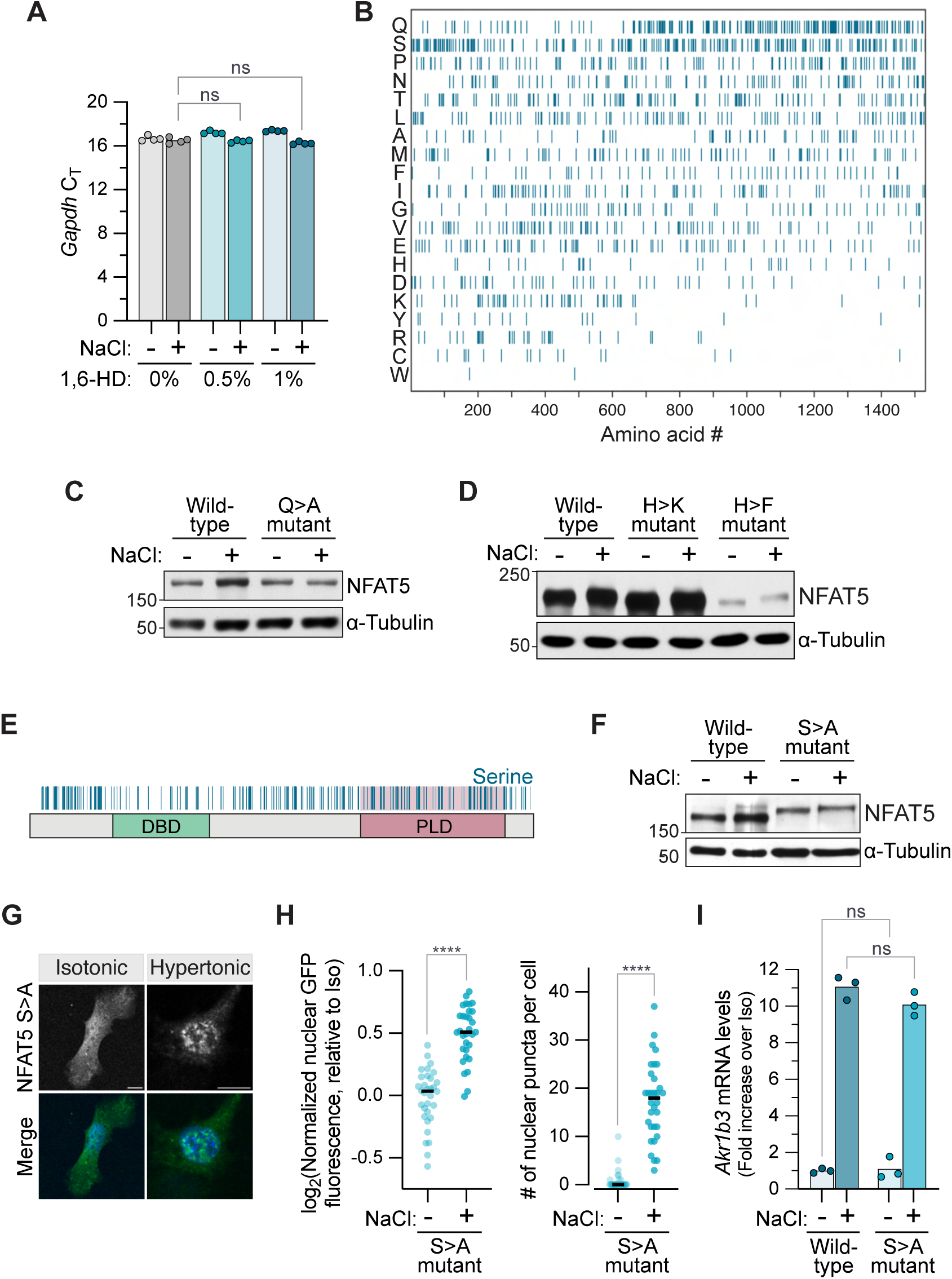
Related to Figure 6. **(A)** Abundance of *Gapdh* mRNA (measured by its C_T_ value in a RT-qPCR assays) after 6 hrs of exposure to 0, 0.5, or 1% 1,6-HD in isotonic or hypertonic (+200 mOsm/L NaCl) media. Compare to the effect of 1,6-HD on *Akr1b3* mRNA abundance in **Figure 6B**. Bars show the mean of 3 measurements. **(B)** Position of each of the 20 a.a. in the NFAT5 linear sequence (oriented along the x-axis). In each row, every occurrence of a single amino acid is marked by a vertical blue line. **(C, D)** Immunoblot comparing the abundances of WT GFP-NFAT5 to the GFP-NFAT5_QA, GFP-NFAT5_HK and GFP-NFAT5_HF mutants. **(E)** Positions of all serine residues in NFAT5 are marked by vertical blue lines. All serine residues in the PLD (highlighted in red) were mutated to alanine residues in the GFP-NFAT5_SA protein. **(F)** Immunoblot comparing abundances of WT GFP-NFAT5 to the GFP-NFAT5_SA mutant. **(G)** Subcellular distribution of GFP-NFAT5_SA stably expressed in *Nfat5*^-/-^ IMCD3 cells after 30 min in hypertonic media (+200 mOsm/L NaCl). Fluorescence signal from NFAT5 is shown alone (top) and merged with DAPI to mark nuclei (bottom). Images of the type shown in (**G**) were used in (**H**) to measure nuclear NFAT5 fluorescence (left) and number of NFAT5 nuclear puncta (right) per cell (*n*>30 cells, with black horizontal line showing the median and each point showing the measurement in one cell). (**I**) Expression of the NFAT5 target gene *Akr1b3* in IMCD3 cells stably expressing WT GFP-NFAT5 or the GFP-NFAT5_SA mutant after 8 hrs in isotonic or hypertonic (+200 mOsm/L NaCl) media. Bars denote the mean of 3 measurements, each shown as points. Scale bars for panel (**G**): 10 μm **Statistics**: For (**A**,**I**), statistical significance was determined by a two-way ANOVA test, Sidak’s multiple comparison. For (**H**) statistical significance was determined by a Kruskal-Wallis test, Dunn’s multiple comparison. P-value symbols are: **** *p*-value<0.0001.

The NFAT5_QA variant is compositionally very different from wild-type NFAT5, having lost its prion-like character. Therefore, we sought to make more subtle mutations that would tune NFAT5 self-associative behavior, using *in vitro* phase separation assays (**Figure 5**) as a guide. We focused on histidine residues because the pKa of its imidazole side chain is known to be sensitive to the ionic strength of the solution.^9,73^ Changes in ionic strength at physiological pH can change the balance between the protonated (positively charged) and deprotonated (neutral) forms of histidine’s side chain. Additionally, the imidazole side chain of histidine has aromatic character, and aromatic a.a. have been shown to function as “stickers” in mediating associative IDR-IDR interactions.^62,64,74,75^ We mutated seven histidine residues within the NFAT5 PLD to lysines (NFAT5_HK) or phenylalanines (NFAT5_HF), mutations designed to weaken or strengthen sticker strength respectively (**Figure 6G**).^76^ *In vitro* phase separation assays were conducted using purified GFP-CTD proteins bearing either the HK or HF mutations (**Figure S6A**). Compared to the wild-type CTD (**Figure 5H**), the phase diagram of GFP-CTD_HK revealed that it had a lower propensity to phase separate, requiring higher NaCl concentrations to form droplets at all protein concentrations tested (**Figures 6H** and **6I**). GFP-CTD_HF formed tangled precipitates of irregular morphology in response to increasing ionic strength, in contrast to the mostly spherical droplets observed with GFP-CTD_HK or GFP-CTD (**Figure 6H**).

Having characterized their self-associative behaviors under purified conditions, we stably expressed GFP tagged full-length NFAT5_HK and NFAT5_HF in *Nfat5*^-/-^ IMCD3 cells to test their abilities to respond to hypertonic stress. The NFAT5_HF mutant formed polymorphic puncta restricted to the cytoplasm even in the absence of hypertonic stress (**Figures 6J**). The apparent lower abundance of NFAT5_HF (as measured by immunoblotting, **Figure S7D**) is likely because it forms aggregates that do not enter polyacrylamide gels. NFAT5_HF failed to accumulate in the nucleus and activate target genes in response to hypertonic stress (**Figures 6K** and **6L**). The abundance of NFAT5_HK was comparable to wild-type NFAT5 (**Figure S7D**) and it accumulated in the nucleus normally in response to hypertonic stress (**Figure 6K**). However, consistent with its reduced self-associative behavior *in vitro* (**Figure 6I**), it formed fewer visible puncta in the nucleus and its transactivation capacity was impaired compared to the wild-type protein (**Figures 6L**). The HF mutations likely enhance or alter self-association in a manner that drives the formation of inactive cytoplasmic aggregates rather than dynamic assemblies or condensates. Thus, the precise nature of the chemical interactions that the NFAT5 CTD makes with itself or other IDRs can tune its transactivation capacity.

Activation of gene transcription in eukaryotes is triggered by sequence-specific transcription factors (TFs) promoting the recruitment or elongation activity of RNA Polymerase II (Pol II) with the help of co­activators like the multi-subunit mediator complex.^63,77^ Recent work suggests that multivalent homotypic and heterotypic IDR-IDR interactions are at the core of these transactivation reactions-- either by promoting the formation of dynamic clusters or co-condensates that include both TFs and co-activators.^68,78–84^ While the physical nature of these transcriptional assemblies and the chemical interactions that drive their formation are debated, our work suggests that transcriptional activation by changes in ionic strength must involve the heterotypic association between the NFAT5 IDR and IDRs from co-activators like the mediator complex. Using a previously established two-color mixing assay, we tested whether the NFAT5 CTD could co-condense with the IDR from the MED1 subunit of the mediator complex (often used as a proxy for the full mediator complex in purified *in vitro* assays).^80,82,85^ At low ionic strength, GFP-CTD remained diffuse and did not show any enrichment in mCherry-MED1_IDR_ condensates (**Figure 7A**). When ionic strength was increased using NaCl, GFP-CTD condensed into droplets that showed almost complete overlap with the mCherry-MED1_IDR_ droplets. The high degree of enrichment of GFP-CTD (and GFP-CTD_HK) in mCherry-MED1_IDR_ condensates is consistent with co-condensation. Interestingly, the aggregates formed by GFP-CTD_HF (which is inactive in cells) were excluded from the mCherry-MED1_IDR_ droplets. Thus, ionic strength promotes IDR-IDR interactions that both drive the self-association of NFAT5 and its association with co-activators, providing a plausible path to the formation of transcriptional assemblies that activate transcription at target genes.

**Figure 7.**
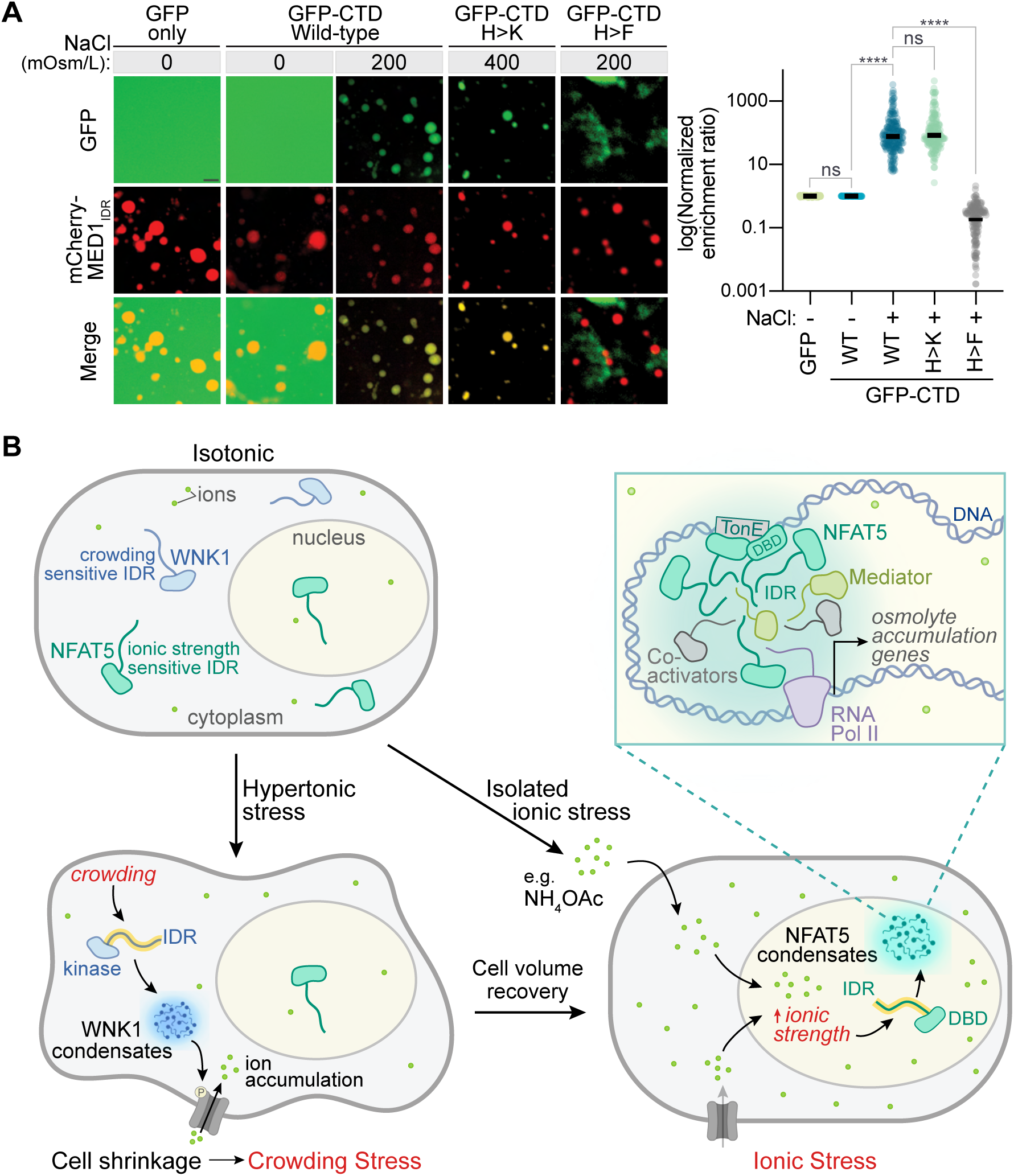
A model for hypertonic and ionic stress adaptation. **(A)** Fluorescence microscopy (left) was used to measure the propensity of GFP, GFP-CTD, GFP-CTD_HK and GFP-CTD_HF (all green) to co-condense with the IDR of MED1 (mCherry-MED1_IDR_, red). All reactions contained 5% Dextran and the indicated concentrations of NaCl. Enrichment ratio (right) of GFP, GFP-CTD, or GFP-CTD mutants in MED1 _IDR_ complex droplets (*n*>50 droplets). Solid horizontal lines denote the median from the individual measurements shown as points. Statistical significance was determined by a Kruskal-Wallis test, Dunn’s multiple comparison. P-value symbol is: **** *p*-value<0.0001. **(B)** Dual IDR-mediated responses to hypertonic and ionic stress in animal cells. The IDRs in WNK1 and NFAT5 each sense different chemical properties of the intracellular environment that are relevant to cell adaptation at short and long timescales. At short time points (min), increase in macromolecular crowding caused by water loss and cell shrinkage is sensed by the WNK1 IDR, triggering a kinase cascade that promotes net ion influx and rapid cell volume recovery (RVI).^41^ However, the rapid recovery of cell volume and intracellular water content comes at the cost of increased intracellular ionic strength, which is incompatible with cell survival over longer time scales (hours). The NFAT5 IDR senses this ionic stress and triggers a transcriptional response that allows replacement of these ions with osmolytes that are non-perturbing to macromolecules and organelles even at high concentrations. We speculate that NFAT5 has evolved to sense and facilitate adaptation to diverse ionic stressors (even those, like NH_4_OAc, that do not cause hypertonic stress).

## DISCUSSION

Even in multicellular animals with systemic neuro hormonal systems, individual cells in tissues critically depend on cell-autonomous osmoregulatory mechanisms to maintain cell volume and intracellular ionic strength homeostasis. The importance of these mechanisms can be seen in cells that function in extreme tissue microenvironments. The urine-concentrating capacity of the kidney depends on the medulla maintaining an interstitial fluid that contains ions at concentrations several-fold higher than their plasma concentrations.^14^ The osmolarity of lymphoid tissues such as the thymus or the spleen can be >30 mOsm/L higher than that of plasma.^86^ Extreme tissue environments are also seen in pathological conditions. Hyperglycemia (seen in diabetes), hypernatremia (caused by unreplaced free water losses or diabetes insipidus), liver failure and kidney failure can all impose chronic hypertonic stress on multiple tissues.^87^ Finally, tumor interstitial fluid has been noted to have markedly elevated K^+^ concentrations (up to 80 mOsm/L above plasma) that influence lymphocyte effector function and exhaustion.^88^

NFAT5 is the master regulator of the cell-autonomous transcriptional response to hypertonic stress in multicellular animals. The complete loss of NFAT5 in the mouse leads to partially penetrant mid-gestational embryonic lethality, with surviving homozygous embryos characterized by massive atrophy of the renal medulla.^89^ Heterozygous embryos display impaired lymphoid proliferation and adaptive immune responses.^86^ NFAT5 null mice and cells completely fail to upregulate osmolyte genes in response to hypertonic stress. These mouse genetic studies show that even without specific stressors, NFAT5 acts as a key guardian of tissues like the renal medulla, likely because of their baseline vulnerability or exposure to hypertonic stress. Intriguingly, variants in *NFAT5* have been identified as quantitative trait loci that impact serum osmolarity and blood pressure in human populations, suggesting a potential role in systemic water and salt balance.^90^

Our work resolves a 25-year-old mystery around the signaling mechanism by which NFAT5 is activated by hypertonic stress. Unlike the yeast HOG pathway, which is structured like a classical signaling system with membrane sensors, kinase transducers and transcriptional effectors, the mammalian hypertonic stress pathway uses NFAT5 both as the sensor and effector. NFAT5 does not directly sense the osmotic pressure gradient across the plasma membrane; instead, we find that it senses the increase in intracellular ionic strength that is a secondary consequence of hypertonic stress. We propose that NFAT5, which is partially localized in the nucleus even under isotonic conditions, is designed to sense ionic strength in the nucleus. Ionic strength-triggered increases in IDR-IDR association between NFAT5 molecules and other components of the transcriptional machinery drive both nuclear NFAT5 accumulation and transcription of its osmo-adaptive target genes.

Hypertonic stress results in two distinct consequences for cells: increased macromolecular crowding at short (minute) timescales and increased ionic strength at long (hour) timescales (**Figure 3A**). Both can disrupt biochemical processes, impair protein folding and promote protein aggregation (often irreversibly).^4,5,40,91^ These distinct chemical changes must require very different sensory mechanisms for their detection. In animal cells, the IDRs of WNK1 and NFAT5 drive the self-association of these proteins into condensate-like assemblies selectively in response to increases in macromolecular crowding and ionic strength, respectively (**Figure 7B**). These environmentally sensitive IDRs are linked to biochemical activities that control the different cellular programs required to resolve these distinct stresses. WNK1 condensation initiates a kinase cascade that promotes Na^+^ influx and inhibits K^+^ efflux, thereby promoting volume recovery and alleviating macromolecular crowding (at the expense of elevated ionic strength).^41^ The NFAT5 IDR drives associative interactions that activate the transcription of osmo-adaptive genes. These observations highlight the exquisite selectivity of IDRs in sensing chemical properties of the intracellular environment, allowing them to initiate responses precisely tailored to restoring homeostasis (**Figure 7B**).^61^

Ionic stress, which is distinct from hypertonic stress, has received scant attention in the literature on stress pathways in animal cells (though it has been investigated in plants). Indeed, we speculate that the function of NFAT5 is not restricted to hypertonic stress responses, but rather NFAT5 maintains intracellular (and intra-nuclear) ionic strength in the optimal window required for the function of most cellular processes. This may be especially important for the nucleus, where protein-nucleic acid interactions are known to be sensitive to ionic strength. By allowing the exchange of excess ions for compatible osmolytes, NFAT5 maintains ionic strength homeostasis. Several processes, such as transepithelial transport, nutrient or waste accumulation, neuronal activity or alterations in transcellular ion fluxes, can change intracellular ionic strength in the absence of elevated extracellular osmolarity. Future studies will investigate how cellular responses to ionic stress are different from previously-described responses to hypertonic stress. Interestingly, intracellular sodium can regulate the material properties of the cell volume regulatory kinase ASK3^67^, suggesting a wider set of ionic strength sensitive pathways. Permeable salts (analogous to NH_4_OAc used in this study) provide a simple and powerful way to impose ionic stress on cells without causing cell shrinkage, water loss and increased macromolecular crowding.

### Limitations of this study

While we have focused on the activation of NFAT5 by hypertonic and ionic stress, our work does not discount a role for other mechanisms (such as phosphorylation) that modulate NFAT5 activity. Indeed, it will be interesting to study how NFAT5 integrates ionic strength signals with signals from other pathways to activate context-or tissue-specific gene expression programs.

The coalescence of NFAT5 into droplets provides evidence that increasing ionic strength can change its chemical properties to promote self-association. These droplets, both *in vitro* and in cells, appear to be formed by phase separation, a segregative density transition. While phase separation propensity is correlated with NFAT5 activity, our data do not prove that phase separation is required for transcription. Indeed, it is possible that dynamic, multivalent clusters of NFAT5 form transcriptionally active hubs in the absence of phase separation.^79^ Broadly, the question of whether multivalent associative clusters or segregative phase separation drives transcriptional activation is an actively debated question for many transcription factors.^63^ How ionic strength increases productive homo-or hetero-typic IDR-IDR interactions to activate transcription is also an important question for future studies. Our mutagenesis experiments point to the importance of histidine residues. The unique pKa of histidine, positioned close to the intracellular pH, may allow it to regulate the conformational ensemble of IDRs in response to changes in the intracellular environment.^9^ We hope that ionic stress-activated transcription emerges as a useful system to investigate these important future questions in the context of NFAT5 and other environmentally responsive transcriptional programs.

## ACKNOWLEDGEMENTS

We would like to thank Dario R. Alessi for generous help with WNK1 pathway constructs and antibodies and Shahar Sukenik for discussion and comments on the manuscript. RR was supported by a grant from the National Institutes of Health (GM118082) and AML by the Intramural Research Program of the National Institutes of Health, National Cancer Institute, Center for Cancer Research. Mengxiao Ma, PhD was supported by an American Cancer Society - Jean Perkins Foundation Postdoctoral Fellowship in Cancer Research, PF- 20-121-01-TBE. The project described was partly supported by Award Number 1S10OD01227601 from the National Center for Research Resources (NCRR).

## AUTHOR CONTRIBUTIONS

RR and HBS conceived the project and HBS first proposed the hypothesis that NFAT5 senses hypertonic stress via a mechanism based on phase separation. HBS established and characterized the reporter system, performed the CRISPR screen (with computational analysis by BBP), and designed and made many key NFAT5 constructs (including mini-NFAT5 and the 7His NFAT5 mutant). CBK designed and performed the haploid genetic screen, with guidance from AML. AML and DT performed the sequencing and computational analysis for the haploid screen. CBK and HBS validated screen hits. CBK designed and performed the experiments in yeast, with advice from MM, and the experiments to distinguish between crowding and ionic strength. PS designed and performed both *in vivo* and *in vitro* phase separation experiments in collaboration with CBK. RR, CBK and PS wrote the manuscript.

## METHODS

## RESOURCE AVAILABILITY

### Lead contact

Further information and requests for resources and reagents should be directed to and will be fulfilled by the lead contact, Rajat Rohatgi.

### Materials availability

All unique reagents generated in this study are available from the Lead Contact and will be provided upon request.

### Data and code availability

The published article contains all datasets generated and analyzed during this study. This paper does not report original code.

## EXPERIMENTAL MODELS AND SUBJECT DETAILS

### Mammalian cell culture and stress treatments

IMCD3 Flp-In cells were cultured in DMEM/F12 (HyClone Dulbecco’s Modified Eagle Medium with L-glutamine and HEPES) supplemented with 10% fetal bovine serum (Atlanta Biologicals), and penicillin (40 U/ml) and streptomycin (40 μg/ml) (Gemini Biosciences). 293FT cells were cultured in DMEM (Dulbecco’s Modified Eagle Medium) containing high glucose (Thermo Scientific) and supplemented with 10% fetal bovine serum, 1 mM sodium pyruvate (Gibco), 2 mM L-Glutamine (Gemino Biosciences), 1x MEM non-essential a.a. solution (Gibco), penicillin (40 U/ml), and streptomycin (40 μg/ml). HAP1 cells were cultured in IMDM (Iscove’s Modified Dulbecco’s Medium supplemented with 10% fetal bovine serum, 2 mM L-glutamine, penicillin (40 U/ml), and streptomycin (40 μg/ml). HAP1 cells were maintained in media containing Deacetyl-baccatin-III (DAB, 2.5 μM) to prevent conversion to diploidy.^95^ All cell lines were kept in a humidified atmosphere containing 5% CO_2_ at 37°C. To impose hypertonic or ionic stress, the indicated amounts of NaCl, sorbitol, urea, NH_4_OAc, were added to growth media (which has a baseline osmolarity of ∼300 mOsm/L). <u>Thus, the total media osmolarity was ∼300 mOsm/L + added osmolarity of NaCl, sorbitol, urea, or NH_4_OAc.</u>

### Mammalian cell lines

Human near haploid cells (HAP1) have been described in our prior publications^22^ and were validated prior to our screens using propidium iodide staining to ensure they have a haploid DNA content. Human 293T and 293 FT cells were obtained from commercial vendors (see Reagent Table) with certificates of authentication and used at low-medium passage (<20) without additional STR profiling. Mouse IMCD3 Flp-I^n^^96^ cells were engineered to contain an FRT site at a unique site in the genome using the Flp-In system from Thermo Fisher, allowing for Flp-mediated stable gene expression from a single, defined genomic locus to readily create sets of isogenic cell lines. IMCD3 Flp-In cells were validated by ensuring they contain a *lacZeo* cassette at the integration locus as recommended by the manufacturer. Cell lines were confirmed to be mycoplasma-negative when cultures were started in the lab and when there was a change in the growth rate or morphology of the cell lines.

#### IMCD3 Flp-In cells and derivatives

The *Nfat5^-/-^* clonal IMCD3 Flp-In cell line (used for the stable, single-locus expression of all NFAT5 variants described in this paper) was generated using CRISPR/Cas9 with a sgRNA targeting the DNA binding domain (Rel-homology domain) of *Nfat5* (**Table S2**). The sgRNA was cloned into pSpCas9(BB)-2A-mCherr^y^^23^ and transiently transfected into WT IMCD3 Flp-In cells using X-tremeGene 9 DNA transfection reagent (Roche). Two days post-transfection, mCherry positive single cells were sorted and assessed by immunoblotting (to measure NFAT5 protein abundance, transcriptional response to hypertonic stress, and sequencing (**Figures 1B** and **S1A**). Clonal *Akap13*^-/-^(**Figure S1E**) and *Wnk1^-/-^* IMCD3 Flp-In cell lines (**Figure 3G**, **S3H**, and **S3N**) were generated using a two-guide system, co-transfecting pSpCas9(BB)-2A-GFP (Addgene plasmid #48138) and pSpCas9(BB)-2A-mCherry expressing gene-specific sgRNAs along with Cas9 and either GFP or mCherry. Cells expressing both fluorescent markers were single-cell sorted using a Sony SH800S Cell Sorter, and edited clones were assessed for genetic knockout via genomic PCR (to ensure excision of the DNA fragment between the two Cas9 cut sites). For the *Wnk1^-/-^* clonal line, genetic knockout was also confirmed by immunoblotting to measure WNK1 protein abundance (**Table S2**).

All NFAT5 variants described in this study were assessed after stable expression at moderate levels from the same single genomic locus in an otherwise isogenic background lacking WT NFAT5 (to prevent the potentially confounding effects of the presence of both WT and mutant NFAT5 in the same cell). To accomplish this goal, all variants were stably expressed in a clonal *Nfat5^-/-^* IMCD3 Flp-In cell line using site-specific recombination into a single site in the genome using the Flp-In system. Briefly, cells at 40-60% confluency in 6- well plates were transfected with a mixture of 2.7 μg pOG44 Flp-Recombinase Expression Vector (Invitrogen), 0.3 μg expression pEF5-FRT-V5_DEST plasmid (Invitrogen) carrying the gene of interest, and XtremeGene 9 (Roche) in 200 μl Opti-MEM Reduced Serum Medium (Gibco). Positive clones were selected for with the addition of 200 μg/ml hygromycin B to the growth medium. Expression of NFAT5 variants was confirmed by immunoblotting to measure the abundance of the protein encoded by the transgene (**Figure S2B**).

To generate a transcriptional reporter for NFAT5 activity (8TonE-GFP), we replaced the seven TCF/LEF binding sites in a previously described reporter for WNT signaling^97^ with eight concatenated copies of the NFAT5-binding TonE site (5-TGGAAAATTAC-3’). The final 8TonE-GFP reporter contains eight TonE sites, a minimal promoter, a gene encoding EGFP and the 5’UTR of the SuperTOPflash WNT reporter. The full 8TonE-GFP cassette was introduced into the unique Flp-In locus of IMCD3 Flp-In cells using site-specific recombination. These reporter cells were subsequently modified for stable expression of an inducible Cas9 (iCas9) for the genome-wide CRISPR screen shown (**Figures 1F** and **1G**). A clonal cell line for the screen was established based on the abundance of nuclear Cas9 protein.

#### HAP1 cells stably expressing the 8TonE-GFP hypertonic stress reporter

293FT cells growing in a 10 cm plate were transfected with pCF567:pLenti-PuroR carrying the 8TonE-GFP reporter cassette mixed with PEI in Opti-MEM. Virus was harvested and then concentrated by ultracentrifugation in sterile 25×89 mm Beckman centrifuge tubes for 1.5 hours at 65,000×*g* at 4°C using a Beckman SW 32 Ti rotor. Pelleted virus was resuspended in 10 ml IMDM (see below for media compositions). 5 ml of resuspended virus was mixed with 5 ml IMDM and polybrene at a final concentration of 4 μg/ml and added to HAP1 cells. 48 hours after transduction, HAP1 cells were treated with 1 μg/ml puromycin to select for cells with a stably integrated 8TonE-GFP cassette. The resulting puromycin-resistant polyclonal population was used to create the Gene-Trap mutant library for genetic screening.

### Yeast culture and treatments for reporter assays

All yeast cells were cultured with shaking at 30°C. For RNA extraction for HOG pathway analysis, W303a (*MATa leu2-3,112 trp1-1 can1-100, ura3-1, ade2-1,* and *his3-11,15*) cells were cultured in yeast extract-peptone-dextrose (YPD) medium supplemented with 0.004% adenine or complete synthetic medium (CSM) supplemented with 2% glucose and 0.004% adenine. For reporter assays and western blot analysis requiring galactose (Gal) induction (**Figure 2D**), saturated overnight cultures of cells in CSM + 2% raffinose, 0.1% glucose, and 0.004% adenine were diluted to an OD_600_ of 0.3 (3.0×10^6^ cells per ml of media) the next morning in CSM + 2% raffinose and 0.004% adenine lacking glucose. Early log-phase cells were then cultured in CSM supplemented with 2% galactose and 0.004% adenine for 2-4 hours before the application of hypertonic stress as described in the figures. 8TonE-*pCYC1*-GFP fluorescence was measured after 4 hours by BD Accuri C6 Flow Cytometer using a 473 nm laser for excitation and a 520/30 nm bandpass filter to collect emitted light.

## METHOD DETAILS

### Western blotting and immunoprecipitation

IMCD3 cells were washed and scraped with ice-cold 1x phosphate buffered saline (PBS). Cell pellets for western blotting were lysed in RIPA buffer (50 mM Tris-HCl, pH 7.4, 100 mM sodium chloride, 1% NP-40, 0.5% sodium deoxycholate, 0.1% SDS, 0.5 mM DTT, 1x SigmaFast protease inhibitor cocktail from Milliore-Sigma, and 1x PhosSTOP phosphatase inhibitor cocktail from Roche). Lysates were cleared by centrifugation at 20,000×*g* for 30 minutes (min) and protein concentrations were determined using the BCA protein Assay (Thermo Fisher). Lysate volumes containing equal total amounts of protein (by mass) were dissolved in 1x NuPAGE Lithium Dodecyl Sulfate (LDS) sample loading buffer and analyzed by SDS-PAGE. The resolved proteins were transferred onto a nitrocellulose membrane using a wet electroblotting system (Bio-Rad) followed by immunoblotting (more detail provided in^22^).

To measure protein abundance (e.g. GFP reporter protein, **Figure S2G**) in W303a yeast cells expressing mini-NFAT5, 3.0×10^7^ of log-phase cells growing in complete synthetic media (CSM) (Sunrise Science) supplemented with 2% galactose were collected by centrifugation, washed with 1 ml of ice-cold water, and resuspended in Y-PER Yeast Protein Extraction Reagent (Thermo Scientific) mixed with 1x SigmaFast protease inhibitor cocktail from Millipore-Sigma. Resuspended cells were mixed with acid-washed glass beads (Sigma Alrich) and lysed by vortexing. Lysates were then cleared at 20,000×*g* for 20 min at 4°C and equal amounts of protein (by mass) from each sample were analyzed by SDS-PAGE and immunoblotting.

For immunoprecipitation (IP) from W303a cells expressing mini-NFAT5 or its variants (dimerization mutant or DNA-binding mutant, **Figure 2G**), 5.0×10^8^ of log-phase cells growing in CSM supplemented with 2% galactose were collected by centrifugation and washed with 10 ml of ice-cold water. Washed cell pellets were then resuspended in ∼1 ml of lysis buffer (20 mM HEPES pH 7.4, 150 mM potassium acetate, 5% glycerol, 1% NP-40, 1x SigmaFast protease inhibitor cocktail, and 0.5 mM DTT), mixed with acid-washed glass beads, and lysed through a 3 minute bead beater cycle at 4°C (MiniBeadBeater-16, Model 607, 3450 RPM, 115 V, BioSpec Products). Samples were spun at 1,000×*g* for 1 min at 4°C to separate the protein sample from the beads and coarse cell debris and then at 20,000×*g* for 20 min at 4°C to obtain a clear lysate for immunoprecipitation. NLS-mRuby3-mini-NFAT5-1D4 variants were immuno-purified via the 1D4 epitope tag by mixing a lysate sample containing 2.5 mg of total protein with Anti-1D4 antibody (The University of British Colubia) covalently conjugated to Protein A Dynabeads (Thermo Fisher Scientific, Invitrogen). Following a 4 hour incubation at 4°C, beads were washed 3 times with a wash buffer (50 mM Tris-HCl, pH 7.4, 100 mM sodium chloride, 1% glycerol). Proteins captured on the anti-1D4 beads were eluted using 2x NuPage LDS sample buffer (95°C, 10 min). Equal volumes of eluates were analyzed by immunoblotting to measure the abundance of 1D4-tagged NFAT45 variants. The abundance of PGK1 was measured in the lysates to ensure that sample inputs for the IPs contained the same amount of total protein.

### Measurement of mRNA abundance by RT-qPCR

RNA was extracted from IMCD3 cells using TRIzol reagent (Life Technologies), and cDNA was synthesized using iScript Reverse Transcription Supermix (Biorad). Real-Time RT-qPCR for mouse *Akr1b3* (aldose reductase, NFAT5 target gene), mouse *Slc5a3* (sodium/myo-inositol cotransporter), mouse *Slc6a12* (sodium-and chloride-dependent betaine transporter) and mouse *Gapdh* (housekeeping gene) was performed on a QuantStudio 5 Real-Time PCR System (Thermo Fisher Scientific) with the following primers -*Akr1b3* (Fwd: 5’-CCTCAGGGAACGTGATACCT-3’ and Rev: 5’-CAATCAGCTTCTCCTGAGTT-3’), *Slc5a3* (Fwd: 5’- GGCAGCAGACATTGCCGTA-3’ and Rev: 5’-AATCGCCACCCAGGTCATAGA-3’), *Slc6a12* (Fwd: 5’- TCTTGGGCTTCATGTCTCAG-3’ and Rev: 5’-GACCTGACTCAGCCACTTCA-3’), and *Gapdh* (Fwd: 5’- AGTGGCAAAGTGGAGATT-3’ and Rev: 5’-GTGGAGTCATACTGGAACA-3’). Reporter GFP transcript levels were assessed using the primers-Fwd: 5’-GACGTAAACGGCCACAAGTT-3’ and Rev: 5’- GAACTTCAGGGTCAGCTTGC-3’. Transcript levels relative to *Gapdh* were calculated using the AACt method.

For yeast mRNA measurements, 4.0×10^7^ of cells were pelleted, resuspended in TRIzol, and lysed using agitation with acid-washed glass beads (5 min, room temperature). After RNA samples were isolated, samples were treated with DNase at 37°C for 1 hour, followed by addition of EDTA and a 10 minute incubation at 65°C to inactivate the DNase. Complementary DNA was synthesized using the iScript Reverse Transcription Supermix. RT-qPCR for *STL1* (Sugar Transporter-like Protein, HOG pathway target gene) and *ACT1* (housekeeping gene) was performed with the following primers -*STL1* (Fwd: 5’-GTTGCGGTATTTCATCAC-3’ and Rev: 5’-CATAGTTGAACTGTTTACC-3’), *ACT1* (Fwd: 5’-GTGTGGGGAAGCGGGTAAGC-3’ and Rev: 5’- GTGGCGGGTAAAGAAGAAAATGGA-3’).^98^ Transcript levels relative to *Act1* were calculated using the AACt method.

### Pooled genome-wide CRISPR/Cas9 screens in IMCD3 cells

Genome-wide CRISPR and haploid screens to identify positive and negative regulators of NFAT5 activity using a fluorescent transcriptional reporter were performed exactly like our previous screens interrogating the Hedgehog^23^ and WNT^22^ signaling pathways.

CRISPR guide RNA library amplification, lentiviral production, functional titer determination and transduction were performed as previously described.^23,99^ To generate a genome-wide mutant cell population for screening, IMCD3 8TonE-GFP iCas9 cells were transduced with the the Brie genome-wide library of sgRNAs,^27^ which targets 19,674 mouse genes with ∼4 sgRNAs each and includes 1,000 non-targeting controls. Each of forty-five T-225 flasks were seeded with 8.0×10^6^ IMCD3 8TonE-GFP iCas9 cells. After cells recovered for eight hours, ∼2.4×10^6^ lentiviral particles mixed with polybrene (2.5 μg/ml) were added to each flask. After the infection was allowed to proceed for 24 hours, successfully transduced cells were selected by supplementing the media with 2 μg/ml puromycin. Screens were scaled so that each guide was represented in at least 1,000 cells in the starting mutant library. For each screen, iCas9 was induced for 5 days using 1 μg/ml doxycycline to initiate genome editing and then the entire mutant cell population was subjected to hypertonic stress (200 mOsm/L NaCl or Sorbitol added to isotonic media in two separate replicates) for 12 hours. A BD FACS Aria Verdi cell sorter was used to isolate cells with the lowest 5% and highest 5% of 8TonE-GFP reporter fluorescence to identify positive and negative regulators respectively. Guide RNA frequencies in the unsorted and sorted populations were determined by deep sequencing followed by analysis using the MAGeCK computational pipeline,^100^ as described in our previous publications.^23^ Screen results can be found in**Table S1** and all Fastq files from NGS have been deposited into the NIH Short Read Archive (SRA) with the BioProject ID PRJNA1015695.

### Insertional mutagenesis screens in human haploid (HAP1) cells

HAP1 8TonE-GFP reporter cells were subjected to pooled insertional mutagenesis using a gene-trap (GT) bearing retrovirus (**Figure S1H**) as described in our previous publications.^22^

#### Retroviral production and concentration

Each of six T-175 flasks were seeded with 1.5×10^7^ 293 FT cells in antibiotic-free media media. When cells reached 80% confluency (∼24 hours), they were transfected with a mixture of retroviral transfer plasmids with the GT in two reading frames (3.3 μg each of pGT-mCherry and pGT+1mCherry), a plasmid carrying packaging genes (4 μg pCMV-Gag-Pol), a plasmid carrying envelope genes (2.6 μg pCMV-VSV-G) and aplasmid to enhance translation (1.7 pg pAdVAntage), all mixed in 450 μl serum-free DMEM containing 45 μl of X-tremeGENE HP DNA transfection reagent. After ∼16 hours, virus-containing media was harvested twice, 8 hours apart. Harvested virus was filtered through a 0.45 μm low-protein binding syringe filter and concentrated by ultracentrifugation in sterile Thinwall, Ultra-Clear, 25×89 mm Beckman centrifuge tubes with a Beckman SW 32 Ti rotor (65,000x*g*, 1.5, 4°C).

#### Insertional mutagenesis of HAP1 cells with GT retrovirus

Each of three T-175 flasks were seeded with 2.0×10^7^ HAP1 8TonE-GFP reporter cells and, after ∼16 hours, infected twice, 8 hours apart, with the GT retrovirus. Concentrated virus was resuspended in 76 ml of IMDM containing 10% fetal bovine serum (FBS) and polybrene (4μg/ml); 25 mls of this suspension was added to each of the three flasks of HAP1 8TonE-GFP reporter cells. Flow cytometry demonstrated that 76.5% of the 8TonE-GFP HAP1 cells had been successfully transduced by a GT retrovirus based on mCherry fluorescence.

#### Selection scheme for HAP1 cells carrying mutations in genes encoding positive and negative regulators

Seven T-175 flasks were each seeded with 2.0×10_7_ cells. The next day, cells (3.0×10_7_) from one plate were collected to map insertions in the control, unsorted parent cell population without the application of any stress or selection. Cells in the remaining six flasks were subjected to hypertonic stress (100 mOsm/L NaCl added to isotonic IMDM media) for six hours later and then FACS was used to isolate cells with the 10% highest GFP reporter fluorescence to identify positive regulators of NFAT5 (**Figure 1F**). To identify negative regulators, cells (in a second screen) were exposed to low-level hypertonic stress (25 mOsm/L NaCl added to isotonic media), followed by sorting to isolate cells with the 10% highest GFP fluorescence. For each sorted population, genomic DNA was extracted from 3.0×10_7_ cells to map retroviral insertions in the selected population. The experimental, computational and statistical methods used to map and compare retroviral insertions in the unsorted and sorted populations has been described in detail previously.^22^ The only difference was that theIGTIOB score used to assess the insertion bias (**Figure 1G**) scored all GT insertions in the selected population, regardless of orientation. Screen results can be found in**Table S1** and all Fastq files have been deposited into the NIH Short Read Archive (SRA) with the BioProject ID PRJNA1015695.

### Validation pipeline for gene hits from CRISPR screen (Figure S1F)

For each candidate gene hit, two separate guides that targeted early exons of all possible transcripts were chosen and individually cloned into px459 (Addgene plasmid #62988). Pooled knockout cell lines were generated by transfection of IMCD3 8TonE-GFP reporter cells with px459 containing a gene specific guide, and cells positive for sgRNA and Cas9 expression were selected with 2 pg/pl puromycin treatment for 3-5 days. Knockout efficiency was determined by sequencing PCR products that encompass the sgRNA cleavage site, and cell populations with >90% knockout efficiency (confirmed using the Synthego ICE CRISPR Analysis Tool, https://www.synthego.com/products/bioinformatics/crispr-analysis) were immediately tested for NFAT5 GFP reporter fluorescence following exposure to hypertonic stress (200 mOsm/L NaCl added to isotonic media for 7-9 hours) by flow cytometry with a BD Accuri C6 Flow Cytometer using a 473 nm laser for excitation and a 520/30 nm bandpass filter to collect emitted light. Two biological replicates were evaluated for each of the two guides specific for each gene (four total independent replicates). All pooled knockout cell populations were compared with cells transfected with a non-targeting control sgRNA.

### Imaging IMCD3 and 293T cells using Immunofluorescence

Cells adherent to coverslips were washed with 1x PBS and then fixed with 4% (w/v) paraformaldehyde (PFA) in PBS. Following 3 washes with PBS for 5 min each, coverslips were either directly mounted on glass slides or processed for further staining. For immunofluorescence staining, cells were permeabilized with blocking buffer (1x PBS, 1% Normal Donkey Serum, 10 mg/ml BSA, 0.1% TritonX-100) for 30 min. Cells were incubated with primary antibodies in blocking buffer for 1-4 hours at room temperature, washed (3x, 5 min each with Wash Buffer containing 1x PBS, 0.1% TritonX-100), and then incubated (1 hour, room temperature) with secondary antibodies in blocking buffer. After 3 additional incubations (5 min each) in Wash Buffer, coverslips were mounted and sealed on glass slides.Images were collected using a Leica TCS SP8 confocal imaging system equipped with a 63x oil immersion objective. All images were taken of cells fixed with PFA unless noted otherwise.

### Image analysis

Image processing for nuclear NFAT5 levels (**Figures 3I, S3E, S3G, S3K, S3M, 6E, 6K and S7H**) was carried out using maximum intensity projection (MIP) images of the acquired *z*-stacks using CellProfiler. For quantification of nuclear NFAT5, first a mask was constructed using the DAPI image (nuclear marker), and then the mask was applied to the corresponding NFAT5 image where the total fluorescence intensity per cell was measured.

NFAT5 puncta number analysis (**Figures 4C, 4I, 5B, 5D, 6E, 6K, S7H**) was carried out using MIP images of the acquired *z*-stacks using the Analyze Particles function of ImageJ. An automated thresholding function was employed for each image before puncta number analysis. For 293T cells, a region of interest was drawn around each cell and the total number of puncta were counted. In case of IMCD3 cells, NFAT5 nuclear puncta numbers were counted for each cell using the DAPI channel as a reference to identify the nuclear boundary.

### Imaging yeast cells (Figure S2H)

Saturated overnight cultures of yeast cells in CSM + 2% raffinose, 0.1% glucose, and 0.004% adenine were diluted to an OD_600_ of 0.3 (3.0×10^6^ cells per ml of media) the next morning in CSM + 2% raffinose and 0.004% adenine lacking glucose. Early log-phase cells were then cultured in CSM supplemented with 2% galactose and 0.004% adenine for 4 hours before microscopy measurements. Cells were stained with Hoechst 33342 (1 μg/ml for 5 min at room temperature) to visualize the nucleus. Cells were mounted in growth medium, and images were collected using an Olympus IX83 epifluorescence microscope equipped with an Orca Fusion scMOS camera using a x100 oil objective (NA 1.45). Hoechst 33342 and mRuby3 were imaged using the DAPI and TRITC channels, respectively. Image analysis and quantification was done using ImageJ.

### Introduction of NFAT5 variants and the NFAT5 transcriptional reporter into yeast (Figure 2D)

W303a cells were transformed with pAG306-8TonE-*pCYC1*-GFP, constructed by replacing the *GPD1* promoter in Addgene plasmid #14140 (pAG306-GPD) with a 8TonE-p*CYC1*-GFP cassette, which was adapted from a yeast transcriptional reporter for calcineurin activity (pAMS366-4xCDRE-GFP-PEST, Addgene plasmid #138658) by replacing the four calcineurin dependent response elements (CDRE’s) with 8 copies of the TonE binding sequence (5-TGGAAAATTAC-3’).^101^ Genomic integration of pAG306-8TonE-*pCYC1*-GFP was done by linearization of the plasmid with the restriction enzyme StuI within the URA3 marker resulting in homology to *ura3-1* and transformed into wild-type W303a cells. Transformed colonies were selected on CSM -Ura plates supplemented with 0.004% adenine and 2% glucose. One colony with the lowest basal GFP expression was used for further experiments. Reporter cells were transformed with pAG305-*pGAL1* plasmids (Addgene plasmid #14137) containing mini-NFAT5 and its variants or other control genes. Genomic integration of the pAG305-*pGAL1* plasmids was done by linearization of the plasmid with the restriction enzyme BstEII within the LEU2 marker resulting in homology to *leu2-3* and transformed into W303a 8TonE-*pCYC1*-GFP reporter cells. Transformed colonies were selected on CSM -Ura -Leu plates supplemented with 0.004% adenine and 2% glucose. Gene integration and protein expression were assessed by PCR and western blotting, respectively.

Mini-NFAT5 (NLS-mRuby3-DBD-MD1-AD2, **Figure 2A**) was constructed by fusing a strong NLS (the Importin-β binding domain from importin-α or IBB^102^ to the Rel-homology DNA binding domain (a.a. 264-543) and two previously described regions (MD1, a.a. 618-821 and AD2, a.a, 1039-1245, from the C-terminal domain of human NFAT5 shown to be important for responsiveness to hypertonic stress.^32^

### Disruption of HOG pathway genes in yeast

Mutant strains were generated in wild-type W303a cells for the sodium measurements using a fluorescent sensor (**Figure S2J**) and in W303a cells expressing the 8TonE-GFP reporter and mini-NFAT5 variants (for **Figure 2K**, **Figures S2I** and **S2K**) by homologous recombination of gene-targeted, PCR products using an established method.^103^ Mutant strains were produced by replacing the complete reading frame of target genes with the *KANMX, HPHMX,* or *TRP1* cassette. Gene deletions were confirmed by PCR amplification of the deleted locus by assessing HOG pathway target gene (*Stl1*, Sugar Transporter-like Protein) induction by RT-qPCR following a 2 hour treatment with 600 mM (1200 mOsm/L) NaCl in CSM containing 0.004% adenine and 2% glucose (**Figure S2I**).

### Intracellular sodium measurements in yeast using a fluorescent sensor (Figure S2J)

W303a cells (wild-type and mutant) cultured overnight in CSM supplemented with 2% glucose and 0.004% adenine were diluted to 2.0×10^6^ cells per ml. After 4 hours, cells were pelleted and resuspended in fresh isotonic media containing the ING-1 AM sodium sensor (ION Biosciences) in 96 well black-bottom plates. The stock solution of the sodium sensor was prepared according to the Manufacturer’s protocol. Briefly, ING-1 AM was dissolved in DMSO and used at a final concentration of 5 ug/ml in isotonic media with 0.02% Pluronic F- 127 (Invitrogen). Cells were then incubated at 30°C for 1 hour with gentle shaking and further grown in isotonic media or media containing 100-600 mM additional NaCl (to apply hypertonic stress) for 30 min. Fluorescence was measured on a Synergy H1 plate reader (BioTek) using an excitation wavelength of 520 nm and emission wavelength of 550 nm. All readings were corrected for background fluorescence, measured from cells incubated without the sodium sensor.

### Measurements of intracellular sodium content in yeast using inductively coupled optical emission spectroscopy (Figure S2K)

Following a previously published protocol,^104^ 5.0×10^8^ of log-phase yeast growing in normal media (CSM with 0.004% adenine and 2% galactose) or media containing additional 400 or 1600 mOsm/L NaCl for 2 hours were collected by centrifugation (8,000×*g* spin for 3 min at 4°C) and washed twice in ice-cold wash buffer containing 1.5 M sorbitol and 20 mM MgCh. Cell pellets were then resuspended in 1 mL 0.2 μm-filtered water, mixed with acid-washed glass beads, vortexed for 3 min and then boiled for 30 min at 95°C with shaking. Following a 1,000×*g* spin to separate the supernatant from the beads and cell debris, samples were cleared using ultracentrifugation with a 100,000×*g* spin for 30 min at 4°C. Cleared samples were then diluted to 5 ml with water, passed through a 0.45 μm filter, and analyzed using an inductively coupled plasma optical emission (ICP) spectrometer (iCAP 6000 Series, Thermo Scientific) through the Environmental Measurements Facility at Stanford University with the following settings: flush pump rate at 50 rpm, analysis pump rate at 50 rpm, pump stabilization time at 5 seconds, RF power at 1150 W, auxiliary gas flow at 0.5 L/minute, and nebulizer gas flow at 0.5 L/minute. Measurements of sodium in samples were calculated based on a standard curve, with NaCl diluted at various known concentrations in filtered water.

### Isotonic shrinkage (Iso-Shrink) of IMCD3 cells (Figure S3C)

The osmolarity of the various solutions used in this protocol was measured using the Fiske Micro-Osmometer (Model 210). The protocol to induce shrinkage of cells without changing intracellular ionic strength was adapted from previous studies.^41,45^ *Nfat5*^-/-^ IMCD3 cells expressing mVenus-NFAT5 or *Wnk1*^-/-^ IMCD3 cells expressing GFP-WNK1 were incubated for 10 min with in a hypotonic medium (osmolarity of 200 mOsm/L containing 1x MEM essential a.a. (Gibco), 1x MEM non-essential a.a. (Gibco), 1x L-glutamine (Gemini Biosciences), 1x MEM vitamin solution (Gibco), 10 mM glucose (Gibco), 50 mM HEPES pH 7.4 (Gibco), 1 mM sodium pyruvate (Gibco), 10% fetal bovine serum (FBS), 2 mM CaCh, 1 mM MgSO_4_, 5 mM KCl, and 30 mM NaCl). During the last 5 min of hypotonic exposure, an RVI inhibitor cocktail (20 μM benzamil, 100 μM bumetanide, 200 μM gadolinium) was added. Hypotonic media was then exchanged for isotonic media (∼300 mOsm/L) containing the same RVI inhibitor cocktail and cells were analyzed after 30 min at 37°. Cells subjected to the isotonic shrinkage protocol were compared to cells treated with hypotonic, isotonic, or hypertonic media all with or without RVI inhibitors.

### Ionic stress imposition on IMCD3 cells using Nystatin (Figure S3I)

We adapted a previous protocol that used nystatin to increase intracellular ionic strength.^55^ Custom isotonic and hypertonic buffers with either high KCl (Iso-KCl, Hyper-KCl) or high NaCl (Iso-NaCl, Hyper-NaCl) were prepared from the same stock solutions. Osmolarities of all buffers were measured using a Fiske Micro-Osmometer (Model 210). All buffers contained 1x MEM essential a.a., 1x MEM non-essential a.a. solution, 1x L-glutamine, 1x vitamin solution, 10 mM glucose, and 10 mM HEPES pH 7.4. Iso-NaCl (286 mOsm/L) contained 5 mM KCl, 122 mM NaCl, 1 mM CaCl_2_, and 1 mM MgCl_2_. Iso-KCl (289 mOsm/L) contained 89 mM KCl, 5 mM NaCl, and 50 mM sucrose. Hyper-NaCl (505 mOsm/L) contained 5 mM KCl, 240 mM NaCl, 1 mM CaCl_2_, and 1 mM MgCl_2_. Hyper-KCl (503 mOsm/L) contained 205 mM KCl, 5 mM NaCl, and 50 mM sucrose.

*Nfat5*^-/-^IMCD3 cells stably expressing FLAG-mVenus-NFAT5 or *Wnk1*^-/-^ IMCD3 cells stably expressing GFP-WNK1 cells were plated on coverslips 1 day before the experiment. Four total treatments (4 different wells of cells) per cell line were compared in this experiment: (1) Isotonic, (2) hypertonic, (3) isotonic+nystatin, and (4) hypertonic+nystatin. The day after plating cells, the Iso-NaCl buffer was added to all wells and after 10 min at 37°C. Next, samples (3) and (4) were treated with ice cold Iso-KCl buffer containing 50 μg/ml nystatin. Following a 10 minute incubation on ice, the media in sample (4) was exchanged to ice cold Hyper-KCl buffer containing 50 μg/ml nystatin. All cells were left to incubate on ice for 20 min. After 20 min, samples (1) and (3) were washed with warm Iso-NaCl and samples (2) and (4) were washed three times with warm Hyper-NaCl to remove the nystatin. After a 30 minute incubation in Hyper-NaCl (samples 2 and 4) or Iso-NaCl (samples 1 and 3) at 37°C, coverslips were washed twice with PBS, and cells were fixed with 4% (w/v) paraformaldehyde (PFA) at room temperature. Instead of treatment with nystatin, the isotonic (sample 1) and hypertonic (sample 2) control cells were kept in either Iso-NaCl or Hyper-NaCl, but were otherwise exposed to the same temperature changes and final buffers as the nystatin-treated cells.

### Measurement of cell height

The cell height was used as a rapid, convenient proxy for cell volume based on previous studies showing that cell height (or thickness) can be used to estimate relative changes in the volume of adherent cells grown in a monolayer.^40,91,105^ CellMask Deep Red Plasma membrane Stain (Thermo Fisher) was used to visualize the plasma membrane to measure cell heights in the xz plane of confocal image stacks. The CellMask stain was added to cells 15 min before beginning all protocols (isotonic shrinkage, nystatin treatments, or salt treatments) and excess stain removed by 3 washes in isotonic media. Cell height was measured from the flat, basal edge of the cell touching the coverslip to the peak height of the cell along the z-axis using ImageJ. Measurements were normalized to the median cell height of the control sample (untreated; samples in isotonic media).

### Measurement of cell volume

IMCD3 cells were plated at a low density in two-well Lab-Tek chambers (500 cells/well) to obtain single separated cells for volume analysis. Cells were labeled with 50 μg/mL Fluorescein diacetate (FDA) in DMEM/F12 medium for 1 hour in a humidified atmosphere with 5% CO_2_ at 37°C. After washing away excess FDA, cells were switched to hypertonic medium (200 mOsm/L NaCl, NH_4_OAc, sorbitol, or urea added to isotonic media) for imaging. All images were acquired on a Leica TCS SP8 confocal imaging system equipped with a 63x oil immersion objective. Fluorescein was imaged using an excitation wavelength of 488 nm and collecting light emitted between 500 and 600 nm. Images of *z* sections of the entire cell were acquired with a fixed step size of 0.5 μm. Individual cell volume was measured using the 3D object counter plugin in Fiji (NIH,Bethesda) after fluorescence intensity thresholding (above background, determined empirically for each image) of *z*-sections.

### Genetically-encoded sensor for intracellular ionic strength

293T cells were plated in 2-well Labtek glass chamber slides (Thermo Scientific) at 10^5^ cells per well. One day after plating, cells were transfected with pcDNA3.1_ionRD (Addgene Plasmid #172931) mixed with PEI in Opti-MEM. The next day, the medium was replaced by FluoroBrite DMEM without phenol red, and cells were imaged directly in the Labtek glass chamber slides. The sensor was excited using a 405 nm laser, and the emission was split into 450-505 nm channel (to image mCerulean3) and 505-797 nm channel (to image mCitrine). The fluorescence intensity of the cells was determined in ImageJ for each channel. The backgrounds for each channel were subtracted and the mCitrine intensity divided by the mCerulean intensity for each cell.

### Endogenous tagging of NFAT5

To generate an IMCD3 clonal cell line with an endogenously tagged *Nfat5* alleles, *Sp* Cas9 protein was complexed with sgRNA (5’-GGGTCGAGCTGCGATGCCCT-3’) and electroporated with the homology-directed repair template plasmid (pUC19-NFAT5/mNeonGreen with ∼700 bp Left & Right Homology Arms flanking mNeonGreen for N-terminal tagging of NFAT5). Production of sgRNA and *Sp* Cas9 were done as described previously.^106^ For electroporation, Cas9-ribonucleoprotein (RNP) complex was prepared by mixing 100 pmol *Sp* Cas9 protein with 120 pmol sgRNA in Cas9 buffer (20 mM HEPES pH 7.4, 150 mM KCl, 1 mM MgCl_2_, 10% glycerol, and 1 mM TCEP) for 10 min. The RNP complex was added to 500,000 low passage IMCD3 cells and electroporated using a NEPA21 Electro-Kinetic transfection system (Bulldog-Bio). 96 hr post-electroporation, mNeonGreen positive single cells (top 0.1% fluorescence intensity) were sorted into a 96-well plate using a FACS (SH800S, Sony). Positive clonal lines were identified by a dual genomic PCR strategy. “In-In” PCR (forward primer complementary within the NFAT5 left homology arm (LHA) (5’- GGAGTCAGTTCTCCACTCCG -3’) and reverse primer complementary within the NFAT5 right homology arm (RHA) (5’- GCGATTCCAGGTCTAGGTCC -3’)) was used to confirm genomic integration of the homology repair template. “In-Out” PCR (forward primer complementary to the genomic region outside the LHA (5’- GGTGTGGTGGGTAACGTGG-3’) and reverse primer complementary within mNeonGreen (5’- GGCCGACCATATCGAAGTCT-3’)) was used to confirm the correct site of integration in the *Nfat5* gene (**Figure S5C**). Expression of the mNeonGreen-NFAT5 fusion protein was confirmed by immunoblotting using an antibody against NFAT5 and the mNeonGreen tag (**Figure S5E**).

### Gal4 DBD luciferase reporter assays (Figure 5E)

293T cells grown in each well of a 96-well plate were transfected with 40 ng of GAL4 upstream activation sequence-driven firefly luciferase expression construct (Addgene plasmid #64125), 40 ng of pGAL4-DBD-MCS-long plasmids (Addgene #145246) directing expression of NFAT5 fragments fused to the GAL4 DNA-binding domain driven by the cytomegalovirus promoter, and 5 ng of herpes simplex virus thymidine kinase (HSV TK) promoter::renilla luciferase plasmids directing constitutive renilla luciferase expression. Cells were cultured for 24 hours, exposed to isotonic or hypertonic media (200 mOsm/L of NaCl or 200 mOsm/L of NH_4_OAC added to isotonic media) and then lysed in Passive Lysis Buffer (25 μl/well, Promega Dual-Luciferase Reporter Assay System) according to the manufacturer’s instructions. Luciferase activities were measured using the Dual-Luciferase Assay System (Promega) in a Synergy H1 microplate reader (BioTek) equipped with an automatic injector. For each sample, NFAT5-GAL4_DBD_ driven firefly luciferase activity was normalized to HSV TK::renilla luciferase activity (to correct for differences in transfection) and the resulting ratio reported in normalized luciferase intensity units.

### Protein Expression and purification

DNA fragments encoding NFAT5 CTD and PLD (and their variants) were cloned into pET-28a plasmid with C-terminal 6xHis-tag either with or without an N-terminal fusion to superfolder GFP (sfGFP) for fluorescence-based phase separation assays. These bacterial expression plasmids were transformed into *E. coli* competent cells (BL21 [DE3] pLysS, MilliporeSigma, Cat #71403) and plated on Luria Broth (LB) agarose plates containing Kanamycin (50 μg/ml) and chloramphenicol (34 μg/ml). A single colony was picked and grown overnight in 10 ml Terrific Broth (TB) containing Kanamycin (50 μg/ml) and chloramphenicol (34 μg/ml) and used to inoculate a 1 L culture of TB. When the culture reached an OD_600_ of 0.4-0.6, protein expression was induced with 0.25 mM IPTG at 37°C for 4 hours. After the addition of 1 mM PMSF and 1 mM EDTA to the cultures, cells were harvested at 4000×*g* for 15 min at 4°C, and flash-frozen in liquid nitrogen.

For protein purification, bacterial pellets were lysed in B-PER Complete Bacterial Protein Extraction Reagent (Thermo Scientific) (5 mL reagent/g of cell pellet) supplemented with 1 mM phenylmethylsulfonyl fluoride (PMSF), 1× protease inhibitor (SigmaFast Protease inhibitor cocktail, EDTA-free; Sigma-Aldrich) and 1 mM EDTA. To facilitate lysis, cells were incubated at room temperature for 15 min with gentle rocking. The CTD and PLD proteins were in insoluble inclusion bodies, which were collected by centrifugation at 20,000×*g* for 30 min at 4°C. The pellets containing inclusion bodies were washed by resuspension using a Dounce homogenizer in 10 pellet volumes in wash buffer (50 mM Tris pH 7.5, 1 mM EDTA, 5 mM DTT, 1 M urea, 1% Triton X-100, 1 mM PMSF and SigmaFast protease inhibitor cocktail) until a clear supernatant was left after pelleting the inclusion bodies. The washed inclusion body pellet was finally suspended in the wash buffer minus the Triton X-100 and urea and collected by centrifugation at 20,000×*g* for 30 min at 4°C. The washed inclusion body pellet was solubilized using a Dounce homogenizer with 1 ml extraction buffer (50 mM Tris pH7.5, 8 M urea, 1 mM PMSF, 1 mM DTT) per gram wet weight of original cells and incubated at room temperature for 1 hour with gentle rocking. The suspension was centrifuged for 1 hour at 100,000×*g* at 4 °C and the supernatant was filtered through a 0.22-μm syringe filter attached to a disposable syringe and loaded on a Superdex 200 16/60 column (GE Healthcare) equilibrated with a low ionic strength gel filtration buffer (20 mM HEPES pH7.4, 1 mM TCEP). Fractions containing NFAT5 CTD and PLD proteins were identified by SDS-PAGE followed by Coomassie Blue staining, pooled and concentrated using Amicon Ultracel 30K filters (Millipore). Proteins were stored at concentrations ≥20 mg/ml on ice and diluted immediately before use. The MED1-IDR tagged with mCherry-tagged (Addgene# 194545) was purified as described previously.^68^

### *In vitro* phase separation assays

Recombinant NFAT5 CTD and PLD fused to sfGFP fusion proteins at varying concentrations were mixed with 5% Dextran (a molecular crowding agent) in droplet buffer (20 mM Na-HEPES pH7.4, 1 mM TCEP) and NaCl (or other salts) were added from a concentrated stock to achieve the indicated osmolarity. The protein solution was incubated for 10 min at room temperature and loaded onto a glass slide with a coverslip. Slides were then imaged with a ×10 dry objective (NA 0.3) on an Olympus IX83 epifluorescence microscope equipped with an Orca Fusion scMOS camera. Droplets containing sfGFP and mCherry were imaged using the FITC and TRITC channels, respectively. Droplets without GFP were visualized using the brightfield channel. Unless indicated, images presented are of droplets settled on the glass coverslip. All assays were performed at room temperature.

To pellet condensates formed by NFAT5 GFP-CTD (**Figure 5G**) droplet reactions (containing 5% Dextran, 70 μM GFP-CTD, 1000 mOsm/L NaCl in droplet buffet) were centrifuged at 20,000xg for 10 min. Supernatant was removed and condensate pellets were resuspended droplet buffer (20 mM Na-HEPES pH7.4, 1 mM TCEP) with 5% dextran (but no added NaCl) to determine if the condensates were reversible.

### Analysis of MED1-NFAT5 co-condensation (Figure 7A)

Equal concentrations (10 μM each) of purified mCherry-MEDI_iDR_ was mixed with purified GFP, GFP-CTD, GFP-CTD H>K or GFP-CTD H> F in droplet buffers with NaCl and 5% dextran. After a 10 minute incubation at room temperature Droplet reactions were imaged as described above in both the mCherry and GFP channels. To calculate enrichment (**Figure 7A**) of the CTD variants in MED1, droplets were defined as a region of interest in FIJI by the mCherry-MED1-IDR channel, and the mean fluorescence signal of the GFP-NFAT5 variants within that droplet was determined. The fluorescence intensity was divided by the background (diffuse) GFP-NFAT5 signal in the image to generate a Cin/out. Enrichment scores were calculated by dividing the Cin/out of the experimental condition by the Cin/out of a control fluorescent protein (GFP).

### Fluorescence recovery after photobleaching (FRAP)

FRAP studies were performed by live confocal fluorescence microscopy using Leica TCS SP8 confocal imaging system. Images were acquired with a 63x oil immersion objective every 600 ms for 1 min. FRAP was performed using the FRAP module of the Leica Application Suite X software with the 488 nm laser at 100% power, 200 μs dwell time. For full-bleach experiments, three repeats for each spot was used for bleaching at maximum laser intensity. This was sufficient to fully bleach most particles. Half-bleach experiments were performed such that droplets were bleached in one hemisphere for 10 frames with a 488 nm laser. Fluorescence recovery was subsequently recorded every 600 ms for a total of 1 min. Image series were imported into FIJI ImageJ for analysis. To calculate the recovery after photobleaching, additional bleaching during image acquisition was corrected using a control non-bleached region over time. To compare multiple experiments, the normalized fluorescence intensity was integrated over all experiments and plotted against time. The pre-bleach signal intensity was set to 100% and the post bleach intensity to 0% for normalization. Data was plotted as percent recovery vs. time in using Prism 10 (GraphPad).

### Super-resolution imaging

Super-resolution three-dimensional-Structured Illumination Microscopy (3D-SIM) images (**Figure 4H**) were collected as Z-stacks (0.125 μm) using a 100× NA 1.40 objective on a DeltaVison OMX Blaze microscopy system, deconvolved and corrected for registration using SoftWoRx. Final assembly of two-dimensional (2D) maximum intensity projections was performed using Fiji.

### Statistical Analysis

Data analysis and visualization were performed in GraphPad Prism 10. Model figures (Figures 1, A, C, and F; 2, D and J; 3, A, B, and F; 5E; 7B; and S1H) were made in Adobe Illustrator 2023 with icons taken from Biorender.com. The predicted human NFAT5 structure was generated using AlphaFold.

The one-way ANOVA test with Sidak’s multiple comparison or the Kruskal-Wallis test with Dunn’s multiple comparison were used to compare three or more groups with one independent variable. A two-way ANOVA test with Sidak’s multiple comparison was used to compare three or more groups with two independent variables. All comparisons shown were prespecified. All experiments were performed at least three different times with similar results. We note that a small sample size (*n*=3) makes it difficult to assess whether the variance between different samples is comparable. Throughout the paper, the *p*-values for the comparisons from GraphPad Prism 10 are denoted on on the graphs according to the following key: \*\*\*\**p*- value<0.0001, \*\*\**p*-value<0.001, \*\**p*-value<0.01, \**p*-value<0.05, and n.s. Non-significant. Replicates: In Figures 1A, 1B, 1C, 1E, 2C, 5E, 6A, 6B, 6F, 6L, S1A, S1E, S2D, S2F, S2H, S2I, S2K, S7A, and S7I, bars and solid horizontal lines denote the mean value from three independent experiments unless otherwise indicated. In Figures 3D, 3E, 3I, 3L, 4I, 5B, 5D, 6E, 6K, 7A, S3B, S3E, S3G, S3K, S3M, S5D, and S7H, solid horizontal lines denote the median value from >3 independent measurements shown as points.

Genome-wide loss-of-function CRISPR screens were performed twice under independent conditions and the duplicates from each screen were analyzed together using the MAGeCK tool (**Figure 1G**). The haploid retroviral mutagenesis screen was performed once. For CRISPR screen validation (**Figure S1F**), two cell lines expressing different sgRNAS against each candidate gene were generated. Cell lines expressing Cas9 and the sgRNA were analyzed two independent times.

## SUPPLEMENTARY TABLES

**Table S1**. Results from CRISPR/Cas9-based and insertional mutagenesis-based loss of function screens.

**Table S2**. Single guide RNA sequences for CRISPR screen validation and clonal knockout cell line generation

